# *Dictyostelium discoideum* flotillin homologues are essential for phagocytosis and participate in plasma membrane recycling and lysosome biogenesis

**DOI:** 10.1101/582049

**Authors:** Cristina Bosmani, Frauke Bach, Florence Leuba, Nabil Hanna, Frédéric Burdet, Marco Pagni, Monica Hagedorn, Thierry Soldati

## Abstract

The metazoan flotillins are lipid rafts residents involved in membrane trafficking and recycling of plasma membrane proteins. *Dictyostelium discoideum*, a social soil amoeba, uses phagocytosis to digest, kill and feed on bacteria. *D. discoideum* possesses three flotillin-like proteins, termed VacA, VacB and the recently identified VacC. All three vacuolins gradually accumulate on postlysosomes and, like flotillins, are strongly associated with membranes and partly with lipid rafts. Vacuolins are absolutely required for uptake of various particles. Their absence impairs particle recognition possibly because of defective recycling of plasma membrane or cortex-associated proteins. In addition, vacuolins are involved in phagolysosome biogenesis, although this does not impact digestion and killing of a wide range of bacteria. Furthermore, vacuolin knockout affects early recruitment of the WASH complex on phagosomes, suggesting that vacuolins may be involved in the WASH-dependent plasma membrane recycling. Altogether, these results indicate that vacuolins act as the functional homologues of flotillins in *D. discoideum*.

## INTRODUCTION

Phagocytosis is the process used by animal phagocytes of the innate immune system to ingest and kill bacterial pathogens and for antigen presentation. The social amoeba, *Dictyostelium discoideum*, which uses this process primarily for nutrition purposes, has been widely used to study the evolutionary conservation of phagocytosis, owing to its experimental versatility and 3R compliance. In addition, the generic endocytic and phagosomal maturation pathways are highly conserved between *D. discoideum* and animals, making it an excellent model to study these mechanisms (Boulais et al., 2010; Dunn et al., 2018).

The particle to be phagocytosed is first bound and recognized by surface adhesion molecules and receptors. This triggers signaling cascades that allow polymerization of actin at the uptake site, ensuring deformation of the plasma membrane and engulfment of the particle in a phagocytic cup (Bozzaro et al., 2008; Dunn et al., 2018). After closure of the phagocytic cup, the particle is found in a compartment called a phagosome, which will mature by a series of fission and fusion events. To allow digestion and killing of bacteria, the compartment needs to become acidic, proteolytic, oxidative and to accumulate toxic levels of certain metals (Cosson and Lima, 2014). To this effect, during maturation, the phagosome, by fusing with lysosomes, acquires the proton pump vacuolar-ATPase (v-ATPase), which acidifies the compartment, followed by delivery of lysosomal enzymes, activated at low pH (Souza et al., 1997; Clarke et al., 2002). In addition, phagocytic cells produce reactive oxygen species (ROS) via the NADPH-oxidase complex and pump toxic levels of metals inside the phagosome (Soldati and Neyrolles, 2012; Dunn et al., 2018; Barisch et al., 2018), rendering the newly-formed phago-lysosome a highly bactericidal environment. In *D. discoideum*, lysosomes eventually mature into neutral and non-degradative compartments, called postlysosomes (PLs), by retrieval of the v-ATPase and, later, of the lysosomal hydrolases. These recycling steps are regulated by the Wiskott–Aldrich syndrome protein and SCAR homologue (WASH) complex, an actin nucleation promoting factor necessary for activation of Arp2/3, an actin-branching complex (Derivery et al., 2009; Carnell et al., 2011; King et al., 2013). Eventually, the PL undergoes exocytosis by fusion with the plasma membrane, in a Ca^2+^-dependent manner (Lima et al., 2012). In addition to phagocytosis, *D. discoideum* uses a non-specific bulk internalization mechanism termed macropinocytosis, which allows uptake of fluid inside a macropinosome that follows a maturation pathway similar to phagosomes (Buckley and King, 2017).

Early during phagosome maturation, plasma membrane proteins that have been engulfed with the particle are sorted away from the degradative pathway and recycled back to the surface. In mammals, several complexes have been implicated in plasma membrane recycling. The retromer is a pentameric complex that includes sorting nexins (SNX) able to sense and induce membrane curvature via their BAR-domain. Thus, the retromer is able to bind to specific cargoes and spatially segregate them in membrane tubules to allow their retrieval (Gallon and Cullen, 2015). The WASH complex is recruited by retromer subunits to endocytic compartments and is suggested to help spatial segregation of cargoes and/or act as a signaling platform owing to its regulation of actin polymerization (Seaman et al., 2013). The WASH complex was shown to participate with another retrieval complex, retriever, in recycling plasma membrane proteins (McNally et al., 2017; Gershlick and Lucas, 2017). Thus, the retriever and retromer complexes, together with WASH, regulate recycling of specific cargoes to the plasma membrane. Another, “slower”, recycling mechanism sorts surface cargoes to a specific tubular juxtanuclear compartment, termed the endocytic recycling compartment (ERC, Maxfield and McGraw, 2004), which harbors the GTPase Rab11. SNX4 is responsible for directing cargoes from endosomes to the Rab11-positive ERC (Traer et al., 2007). In *D. discoideum*, recycling of plasma membrane proteins from macropinosomes and phagosomes has recently been shown to require the WASH complex (Buckley et al., 2016). Furthermore, receptors delivering lysosomal enzymes to endosomes also need to travel back to the *trans*-Golgi network, to be recycled for several maturation cycles. This transport, termed retrograde trafficking, is regulated by the retromer complex in animals (Burd and Cullen, 2014).

Flotillin-1 and flotillin-2 are lipid rafts proteins conserved throughout evolution in Metazoans (Morrow and Parton, 2005; Otto and Nichols, 2011). They are composed of a PHB domain (prohibitin homology domain) at the N-terminus, responsible for their insertion into the cytoplasmic leaflet of membranes (Morrow et al., 2002). This domain is characterized by hydrophobic stretches and acylation sites, which are required for membrane binding (Morrow et al., 2002; Neumann-Giesen et al., 2004). At the C-terminus lies the so-called flotillin domain, which contains coiled-coil regions, necessary for protein-protein interactions and hetero- and homo-tetramerization (Solis et al., 2007). Flotillins are found in specific detergent-resistant lipid microdomains at the plasma membrane and at endosomal and phagocytic compartments (Dermine et al., 2001; Neumann-Giesen et al., 2004; Liu et al., 2005). At the plasma membrane, flotillins are thought to function as signaling platforms involved in axon regeneration and glucose uptake, among other functions (Baumann et al., 2000; Stuermer, 2011a). In endocytic compartments, flotillins have been shown to participate in trafficking of several cargoes across multiple cell types (Stuermer, 2011b; Meister and Tikkanen, 2014). Recently, flotillin-1 was shown to bind to the endosomal sorting complex required for transport (ESCRT) −0 and −1 and proposed to be involved in transferring cargoes destined for degradation from ESCRT-0 to ESCRT-1 (Meister et al., 2017). On the other hand, the surface transferrin receptor (TfR) and E-cadherin require flotillins to be recycled to the plasma membrane through the ERC in a Rab11a- and SNX4-dependent manner (Solis et al., 2013). Moreover, flotillins interact directly with Rab11a at the ERC in epithelial cells as well as neurons, suggesting that their function in plasma membrane recycling may be conserved in multiple cell types (Bodrikov et al., 2017a).

*D. discoideum* has three flotillin-like proteins, called vacuolins A, B and C. These proteins are highly similar and share a structure reminiscent of the one of flotillins, with a PHB-like domain and coiled-coil regions (Wienke et al., 2006). The coiled-coil regions at the C-terminus mediate oligomerization, and it was shown that VacA and VacB can oligomerize with themselves as well as with each other (Wienke et al., 2006). The PHB domain has also been implicated in membrane targeting, although how vacuolins are associated with membranes has not been investigated yet. Vacuolins bind membranes in a patchy distribution, thereby defining specific microdomains (Rauchenberger et al., 1997). VacA and VacB have been previously described as PL markers in *D. discoideum* AX2-214 background (Rauchenberger et al., 1997; Jenne et al., 1998). However VacC, which was only recently identified after the sequencing and annotation of the genome (Eichinger et al., 2005), has not been studied so far. *vacA* and *vacB* knock-out (KO) mutants were generated in the AX2-214 background, revealing that VacB is involved in PL biogenesis. In fact, in the absence of VacB, cells had a delayed lysosomal reneutralization phase, a longer transit time of fluid phase and delayed exocytosis (Jenne et al., 1998). In that study, no phenotype was observed for *vacA-* cells, suggesting a non-redundant role for both vacuolins (Jenne et al., 1998). In addition, VacB was proposed as a negative regulator of PL fusion, as PLs in *vacB-* cells were enlarged (Jenne et al., 1998; Drengk et al., 2003). Together, these results indicated a role for VacB in PL biogenesis.

We sought to characterize the newly identified vacuolin, VacC, and, in the light of their potential homo- and heteromerisation, revisit the localization, biochemical behavior and role of all three vacuolins. For this purpose, we have generated overexpressing and chromosomally-tagged GFP-fusions of each vacuolin, as well as single and multiple KOs. In this study, we bring further evidence that vacuolins are the functional homologues of flotillins in *D. discoideum*, and show that they are involved in phagocytosis, plasma membrane recycling and phagosomal maturation.

## RESULTS

### The three *D. discoideum* vacuolins gradually accumulate on PLs

To characterize the localization of VacC compared to the other two vacuolins, we inserted GFP in-frame at the C-terminus of each vacuolin gene in wt cells (Vac-GFP KI, Materials and Methods). Cells were immunostained for GFP, the A subunit of the v-ATPase, VatA, which resides in lysosomes and the contractile vacuole (CV), or p80, a predicted copper transporter which accumulates in PLs (**Fig. 1A**). As previously described (Rauchenberger et al., 1997; Jenne et al., 1998), VacA and VacB were present at PL compartments in a patchy distribution, suggesting that they are present in specific microdomains. Similarly, VacC colocalized with p80 at PLs and was not found at lysosomes with VatA (**Fig. 1A-B**). Localization of endogenous (endo) VacA and VacB with newly-produced specific recombinant nanobodies (**Fig. S1**, Materials and Methods) produced similar results (**Fig. 1C**).

**Figure 1:**
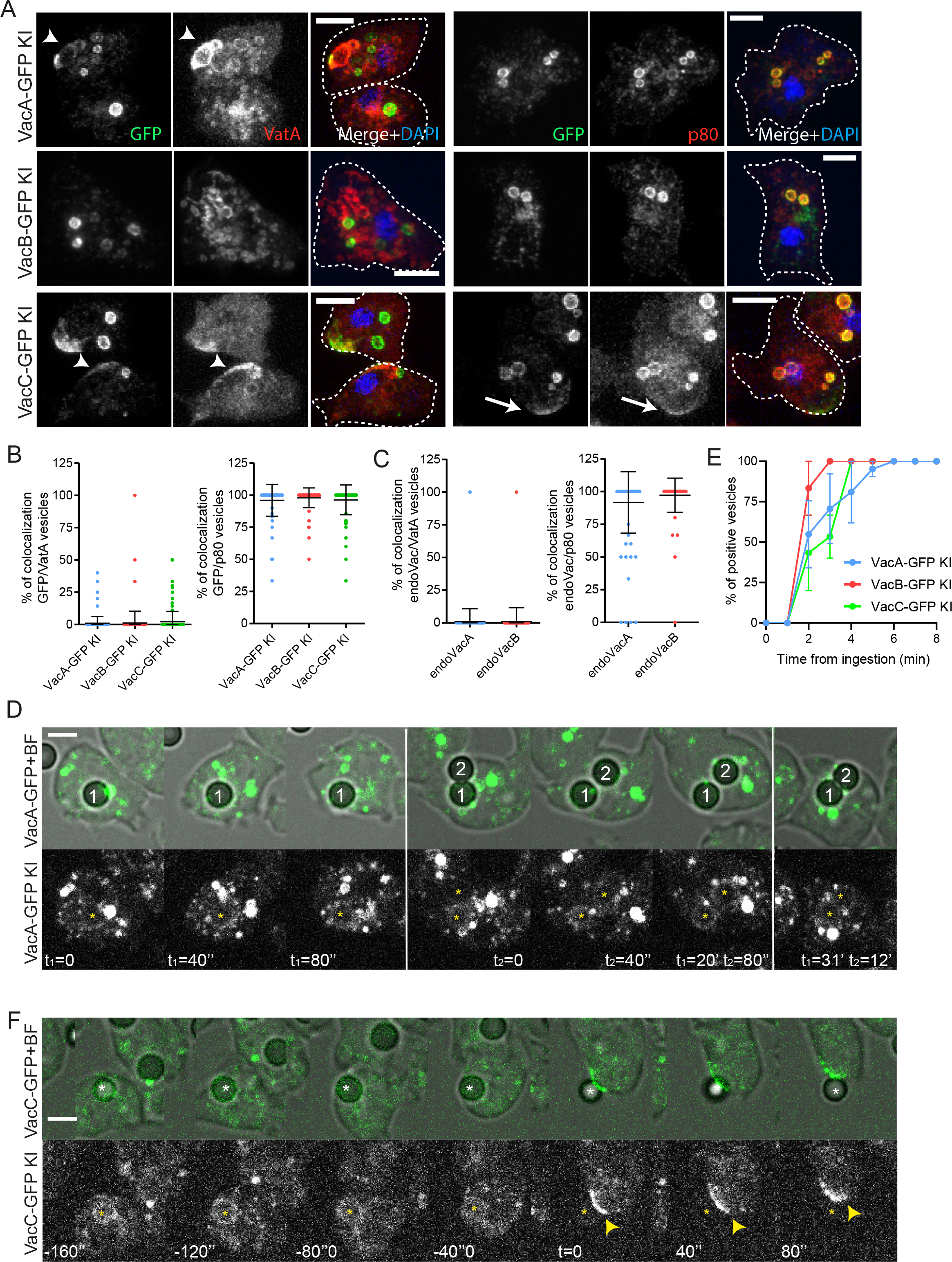
Localization of *D. discoideum* vacuolins. **A-B.** The three vacuolins colocalize with p80 at postlysosomes. **A.** AX2(Ka) cells with GFP knocked-in (KI) at the C-terminus of VacA, VacB or VacC were fixed and immunostained for GFP (green), VatA (red, left panel) or p80 (red, right panel), as well as DAPI (blue). Maximum projections are shown. Arrows, exocytic patches (vac- and p80-positive); arrowheads, CV (vac- and VatA-positive) Scale bars, 5 μm. **B.** Quantification of A. The percentage of colocalization was calculated by counting the proportion of GFP-positive vesicles that were VatA- or p80-positive for each cell. VatA positive CV structures were not quantified. Each dot represents one cell (mean ±sd, n≥200 cells, N=2). **C.** Endogenous VacA and VacB also accumulated at postlysosomes. Wt cells were fixed and immunostained for VacA or VacB and VatA or p80. Quantification was performed as in B. Each dot represents one cell (mean ±sd, n≥100 cells, N=1). **D-E.** Vacuolins are present early at phagosomes. **D.** VacA-GFP KI cells were spinoculated with latex beads and imaged by time-lapse microscopy. Representative Brightfield (BF) and VacA-GFP images are shown. Time from ingestion of the two beads is indicated; asterisks label the phagocytosed beads; scale bar, 5 μm. **E.** Quantification of D. The percentage of vacuolin-positive beads was quantified at the indicated time points (mean ±s.e.m., N=3 for VacA and N=2 for VacB and VacC). **F.** Representative images of a VacC-GFP-positive exocytic patch at the plasma membrane after exocytosis of a bead. Time indicated in seconds, with t=0 set at the exocytosis; scale bar, 5 μm.

Surprisingly, while overexpressed VacA- and VacB-GFP (OE) showed a similar localization as the KIs, overexpressed VacC-GFP exhibited a more heterogenous localization, as it mostly colocalized with VatA-positive lysosomal vesicles (75%) and only partially (25%) with p80 compartments (**Fig. S2A-B**). Further characterization of Vac-GFP OE strains revealed that cells overexpressing vacuolins, in contrast to the KIs, had a larger number of p80-positive compartments compared with wild-type (wt) cells (**Fig. S2C**). In addition, the number of GFP-positive, i.e. vacuolin-positive, vesicles was also increased in overexpressing cell lines with an average of over ten vesicles per cell, compared to three to four vesicles in KIs (**Fig. S2D**). Similar to the KIs, the anti-VacA or anti-VacB nanobodies stained three to four vesicles per cell (**Fig. S2E**). These results are in agreement with the number of vesicles revealed by the 221-1-1 anti-vacuolin antibody (renamed here pan-vacuolin, see Materials and Methods, Rauchenberger et al., 1997). We conclude that overexpression of vacuolins may be detrimental to the cell and therefore not suitable to study the physiological localization and functions of vacuolins.

To study more in detail the dynamic of recruitment of all three vacuolins to phagosomes, Vac-GFP KI cell lines were monitored by live microscopy during phagocytosis (**Fig. 1D-E**). Interestingly, all three vacuolins were present on phagosomes as early as two to four minutes after closure. In fact, phagosomes presented clear, although faint, vacuolin-positive patches as early as two minutes after closure. In addition, phagosomes were surrounded by highly dynamic and vacuolin-rich vesicles. Moreover, we were able to observe vacuolin patches at the plasma membrane (**Fig. 1F**), that resulted from exocytic fusion of the bead-containing phagosome (BCP), as previously hypothesized (Jenne et al., 1998). Together, these results show that all three vacuolins, which are highly similar (**Fig. S1F**), are recruited very early to phagosomes and steadily accumulate on PLs until exocytosis.

### Vacuolins partially partition into detergent resistant membranes

Flotillins are well established markers of lipid rafts at the plasma membrane and phagosomes, and are tightly associated with membranes and resistant to alkaline extraction (Bickel et al., 1997; Dermine et al., 2001; Morrow et al., 2002). Given the similarity in the protein domains of flotillins with *D. discoideum* vacuolins (**Fig. S1F**), we investigated whether vacuolins biochemically behave like their putative mammalian homologues. To test vacuolins’ membrane association, Vac-GFP KIs were homogenized and ultracentrifuged to separate the cytosolic (Cyto) from the membrane-bound (MB) fraction, followed by alkaline extraction and western blotting (**Fig. 2A**). The integral membrane protein, mitochondrial porin, was exclusively found in the membrane fraction. Both the endogenous and Vac-GFP KIs were found exclusively associated with membranes. In addition, this association was quite strong, as alkaline pH only slightly extracted vacuolins from the MB fraction. To better characterize the interaction with the membrane, several extraction methods were applied on cells overexpressing VacB-GFP (**Fig. 2B**). Treatment with urea was not sufficient to extract vacuolins from the MB fractions, but high salt concentration and alkaline pH partially extracted vacuolins, but not mitochondrial porin, as expected.

**Figure 2:**
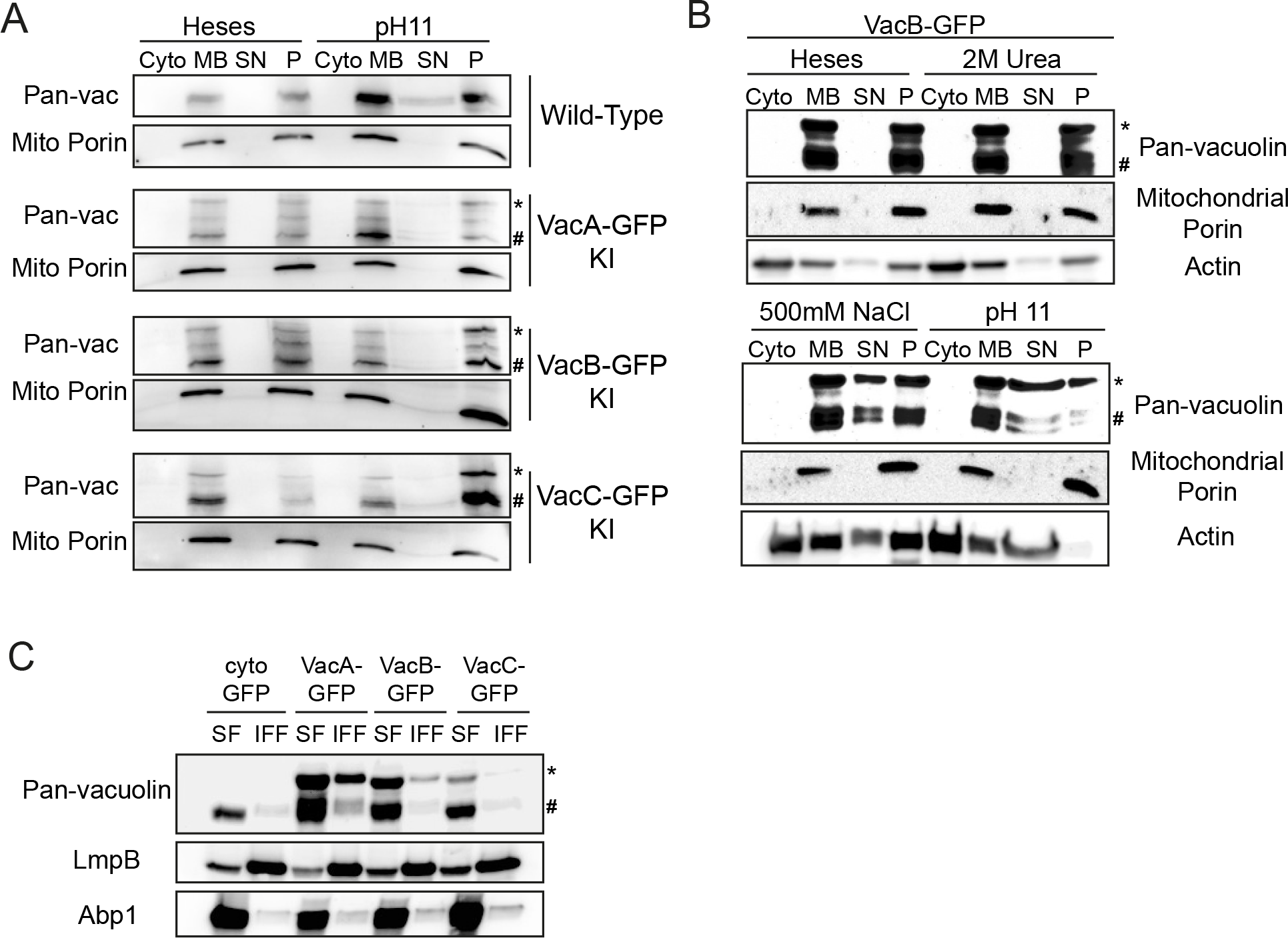
Vacuolins are tightly associated with membranes and partially partition in DRMs. **A-B.** Vacuolins are tightly associated with membranes. **A.** Vac-GFP KIs and wt cells were homogenized and ultracentrifuged to separate cytosolic (Cyto) and membrane (MB) fractions. MB fractions were then treated with Heses or sodium carbonate pH 11 and separated into supernatant (SN) and pellet (P) fractions by ultracentrifugation. Similar amounts of each fraction were further analyzed by western blotting with the indicated antibodies (representative blots, N=2). **B.** Wt cells overexpressing VacB-GFP were treated as in A., membranes were also treated with high concentrations of urea or NaCl (representative blots, N=2). **C.** Vacuolins are partially associated with detergent resistant membranes (DRMs). Wt cells overexpressing cytosolic GFP, VacA-GFP, VacB-GFP or VacC-GFP were lysed in cold Triton X-100. The Triton-soluble fraction (SF) was recovered and the Triton-insoluble floating fraction (IFF) containing the DRMs was isolated after flotation on a sucrose gradient. Fractions were analyzed by western blotting with the indicated antibodies (representative blots, N=2). *, vacuolin-GFP bands (~100kDa); #, endogenous vacuolins (~68kDa).

Proteins that form detergent resistant microdomains, such as flotillins or caveolins, are known to resist to alkaline extraction, despite their lack of transmembrane domains (Schlegel and Lisanti, 2000; Morrow et al., 2002). To test whether vacuolins are found in detergent-resistant membranes (DRMs), we lysed Vac-GFP OE strains with cold Triton X-100 and separated the Triton-soluble (SF) from the insoluble floating fraction (IFF), containing the DRMs, followed by western blot (**Fig. 2C**). LmpB, a plasma membrane CD36-like glycoprotein (Sattler et al., 2018), was enriched in the IFF, as previously described (Harris et al., 2001a, 2001b), while the actin-binding protein, Abp1, was excluded from DRMs. Although vacuolins were mostly soluble in cold Triton X-100, a small fraction of all three Vac-GFP, as well as the endogenous proteins, was found associated with DRMs (**Fig. 2C**). We conclude that, like mammalian flotillins, vacuolins tightly associate with membranes, but are only partially inserted into detergent-resistant microdomains.

### The reported vacB-strain is not a single gene knock-out

In order to better characterize the role of vacuolins, we first resorted to use the previously published gene disruption mutants *vacA-* and *vacB-* in the AX2-214 background (Jenne et al., 1998). However, during our studies we realized that the *vacB-* mutant was not a single KO. To our surprise, while in *vacA-* only the central portion of *vacA* was not amplified by PCR on genomic DNA, in the *vacB-* mutant the central portions of both *vacB* and *vacC* were missing (**Fig. S3A**). RNA sequencing of both *vacA-* and *vacB-* mutants confirmed these results, and revealed that apart from *vacB* and *vacC*, no mRNA was detected over the neighboring *slob2* gene in *vacB-* mutants (**Fig. S3B**). Moreover, reads were missing from the end of the *cax1* gene, suggesting that the deletion in the *vacB-* mutant extended from the central region of *cax1* until the *vacB* gene. For these reasons, we decided to generate new isogenic vacuolin KO mutants, in the AX2(Ka) background, by gene deletion and insertion of an antibiotic resistance cassette (Materials and Methods). Single KO mutants of each vacuolin, as well as a double *vacB/vacC* and a triple vacuolin mutants were generated. These were tested by PCR on genomic DNA, qRT-PCR, western blotting as well as by RNAseq (Materials and Methods and **Fig. S3C**), to ensure that only the intended gene(s) had been disrupted. We therefore used these mutants to completely revisit the role of vacuolins in *D. discoideum*.

### Vacuolins are required for particle uptake

Given that vacuolins are recruited early and are present throughout the phagosomal maturation, we investigated their role in uptake and phagosomal biogenesis. We first tested whether absence of one, two or three vacuolins impaired phagocytic uptake. Cells were incubated in the presence of various-sized fluorescent beads (**Fig. 3A**), GFP-expressing Gram-negative *Klebsiella pneumoniae* and pathogenic *Mycobacterium marinum* (**Figs. 3B-C**). Uptake was measured by following accumulation of fluorescence over time by flow cytometry. All vacuolin KO mutants were severely impaired in uptake of every particle tested, with as much as 90% decreased internalization. This defect was not dependent on the size (**Fig. 3A**) nor on the nature of the particle (**Figs. 3B-C**). Interestingly, vacuolin KO mutants were slightly less impaired in uptake of *K. pneumoniae*, with about 75% reduction (**Fig. 3B**), compared to 95% reduction for beads or *M. marinum* (**Figs. 3A and C**).

**Figure 3:**
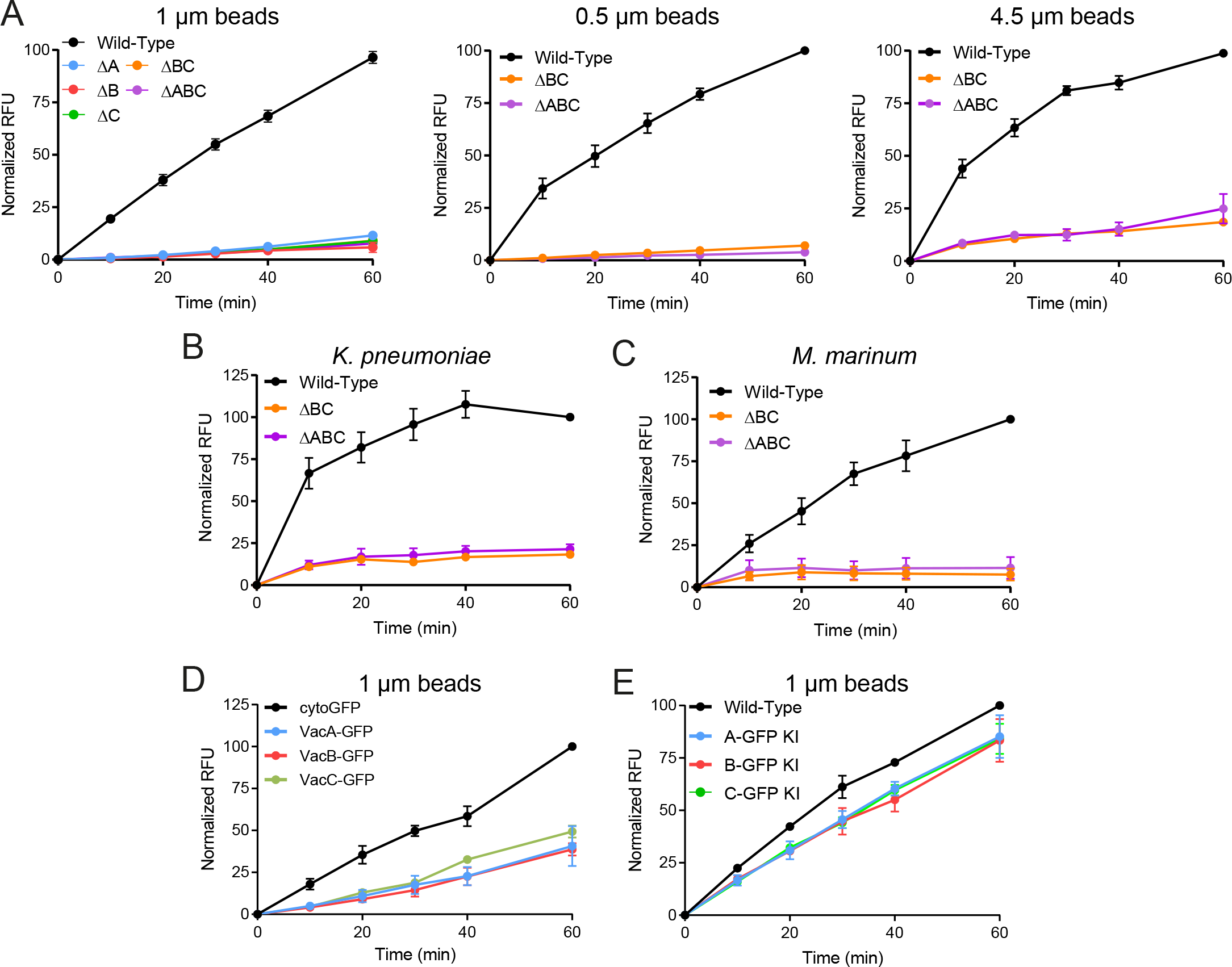
Absence of vacuolins impairs phagocytosis of various particles. **A-C.** Wt and vacuolin single, double or triple mutant cells were shaken for 1 hour in the presence of fluorescent 1 μm, 0.5 μm or 4.5 μm YG-beads (**A.**), GFP-expressing *K. pneumoniae* (**B.**) or *M. marinum* (**C.**). Cells were collected at the indicated time points and fluorescence measured by flow cytometry. Graphs depict Relative Fluorescence Units (RFU) normalized to wt cells at 60 min (mean ±s.e.m., N=3). **D.** Same as in A. with cells overexpressing cytosolic GFP, VacA-GFP, VacB-GFP or VacC-GFP with 1 μm beads. (N=2). **E.** Same as in A., with VacA-GFP, VacB-GFP, VacC-GFP KIs and 1 μm beads. (N=2).

Overexpression of the three Vac-GFP in wt cells leads to an increased number of PL compartments and a different localization for VacC (**Fig. S1**). To test whether overexpression also affected uptake, Vac-GFP OE cells were also assayed, revealing a 30 % reduction in uptake of 1 μm beads (**Fig. 3D**). In contrast, expression of chromosomally GFP-tagged vacuolins did not affect uptake (**Fig. 3E**). We conclude that both absence and overexpression of vacuolins severely impairs uptake of various particles, suggesting that the absolute quantity of each vacuolin as well as their stoichiometry may be important for their function.

### Absence of vacuolins impairs particle recognition

Vacuolins, unlike flotillins, are present only transiently at the plasma membrane, upon PL exocytosis, suggesting that their role in phagocytic uptake is indirect. Inability to take up particles may be explained by impaired motility towards the particle, recognition or defects in adhesion. In addition, actin-dependent rearrangements responsible for the formation of the phagocytic cup, phagosomal closure or signaling to initiate uptake may be affected as well. To investigate how absence of vacuolins impaired uptake, we subjected the ΔABC cells to various assays to test which phagocytosis step may be affected.

To test whether formation of the phagocytic cup and its closure were impaired, wt and ΔABC cells were transfected with the phosphatidyl-(3,4,5)-triphosphate (PI(3,4,5)P_3_) reporter PH_CRAC_-GFP that is enriched at phagocytic cups and macropinosomes in *D. discoideum* (Parent et al., 1998; Buckley et al., 2016). To observe PH_CRAC_-GFP dynamics upon phagocytosis, cells were imaged by time-lapse microscopy in the presence of TRITC-labelled yeasts (**Fig. 4A**). We first observed that the morphology of the forming phagosome around the particle was often different in ΔABC cells compared with wt. Notably, in wt cells, the phagosomal membrane was tightly apposed to the particle, whereas in ΔABC cells, particles were more frequently taken up in structures that resembled widely open macropinosomes (**Fig. 4A**). Accumulation of PI(3,4,5)P_3_ at the site of the forming phagosome is required for efficient closure of the phagocytic cup in mammalian cells (Dewitt et al., 2006). To determine whether phagosomal closure was affected in ΔABC cells, we measured the time from first appearance of the PI(3,4,5)P_3_ patch until its disappearance, and thus phagosomal closure (**Fig. 4B**). No significant difference was observed between wt and ΔABC cells, suggesting that PIP-dependent signaling and phagosomal closure are not affected by vacuolin absence.

**Figure 4:**
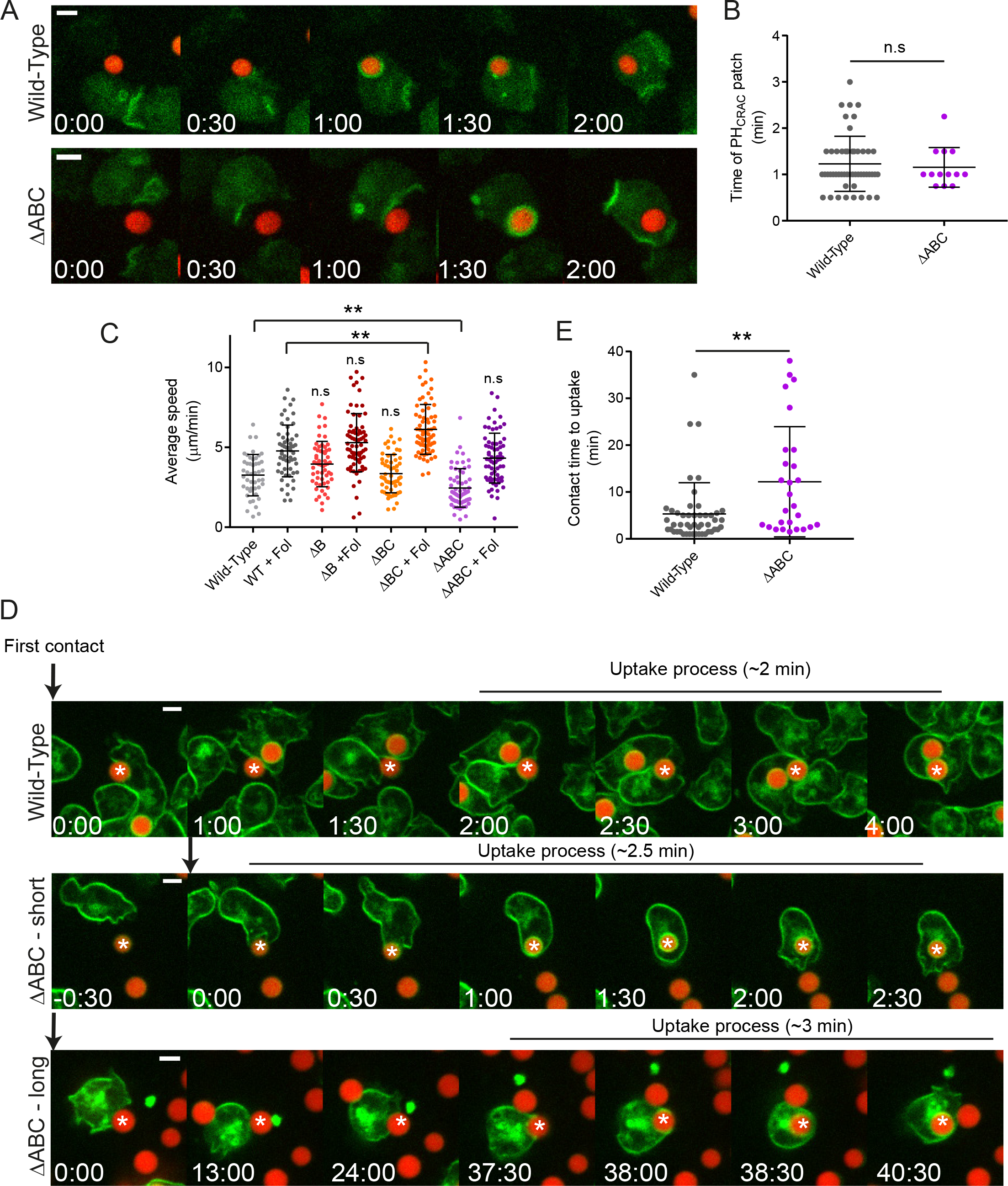
Absence of vacuolins impairs particle recognition. **A-B.** Phagosome closure is not affected in triple vacuolin mutant cells. **A. Wt** or ΔABC cells overexpressing PH_CRAC_-GFP (green) were plated with TRITC-labelled yeasts (red) and imaged by time-lapse microscopy. A representative example is shown for each cell line. Time is indicated in minutes and seconds. Scale bars, 5 μm. **B.** Quantification of A. The time from the appearance of the PH_CRAC_-GFP patch until its disappearance, and thus closure of the phagosome, was measured. Each dot represents one event (mean ±sd, n=50 events for wt, n=13 for ΔABC, N=3, Mann-Whitney test). **C.** Absence of vacuolins only slightly impacts motility. Wt or mutant cells were plated at low confluency with or without folate (Fol) and imaged by high-content microscopy every 30 seconds. Cells were tracked using the MetaXpress software and average speed was calculated. Each dot represents one cell. For statistics, mutant cells were compared with the corresponding wt condition (± folate) (mean ±sd, n≥70 cells, N=1, one-way ANOVA, Kruskal-Wallis test, **p≤0.01). **D-E.** ΔABC cells take longer to initiate phagocytosis. **D.** Wt or ΔABC cells were stained with the fluorescent membrane dye FM4-64 (green), plated in the presence of fluorescent beads (red) and imaged by time-lapse microscopy. Snapshots of representative examples are shown and time is indicated in minutes and seconds. Scale bar, 5 μm. Asterisk points to the phagocytosed bead. **E.** Quantification of C. The time from the first contact with the bead until its internalization was measured. Each dot represents one event (mean ±sd, n=48 for wt, n=27 for ΔABC, N=3, Mann-Whitney test, **p≤0.01).

To determine whether vacuolin KO cells are less motile, and thus less capable of moving towards particles, we assayed their random motility and velocity in phosphate buffer, in the presence or absence of the chemoattractant and motility stimulator folate (**Fig. 3C**; (Lima et al., 2014)). ΔABC cells moved slightly, though significantly, more slowly than wt cells, however, upon stimulation with folate, their random motility was the same as wt. In addition, the motility of ΔBC cells was slightly higher than wt upon folate stimulation. Nevertheless, we conclude that these results do not reveal a major actin-dependent motility defect that may explain the severe uptake phenotype observed with vacuolin KOs.

Finally, we observed that, when imaged by time-lapse microscopy, vacuolin KO mutants were able to take up particles, although less frequently than wt cells and often after prolonged contact. Therefore, we wondered whether absence of vacuolins perturbed particle recognition and adhesion. To test this hypothesis, the plasma membrane of wt and ΔABC cells was labelled with the lipophilic dye, FM4-64, to follow formation of the phagosome after uptake of fluorescent beads. We measured the time from the first contact with the particle until phagosomal closure, as an estimation of the time it took to recognize the particle (**Fig. 4D**). Wt cells took on average 2 to 5 minutes to engulf completely the particle after first contact (**Fig. 4E**). On the other hand, ΔABC displayed a more heterogenous phenotype, with cells taking as short a time as wt to engulf particles, and others as long as 30 to 40 minutes. In conclusion, these results suggest that absence of vacuolins may indirectly impair initial recognition and/or adhesion to the particle and initiation of phagosome formation.

### Vacuolins are involved in plasma membrane protein recycling

Flotillins have been recently shown to localize to the Rab11a-positive ERC, and, in concert with Rab11a and SNX4, participate in the recycling of the plasma membrane proteins TfR and E-cadherin (Solis et al., 2013; Bodrikov et al., 2017b). Given the similarity between mammalian flotillins and vacuolins, we wondered whether they may be also involved in plasma membrane recycling. In fact, we reasoned that a defect in recycling may affect the presence of adhesion and receptor molecules at the cell surface and thus uptake. To verify this hypothesis, we biotinylated the cell surface of wt and ΔABC cells and purified phagosomes at different maturation stages, to assay the dynamics of plasma membrane proteins retrieval from phagosomes (**Figs. 5A-B**). Latex bead-containing phagosomes were isolated by flotation on sucrose gradients and analyzed by western blot. Streptavidin-HRP was used to reveal all surface proteins present in the phagosomes. The presence of the β-integrin-like protein SibA and glycoprotein LmpB, two known plasma membrane proteins (Janssen et al., 2001; Cornillon et al., 2006; Sattler et al., 2018), were used as control. Importantly, latex beads were first adsorbed on cells in the cold before induction of phagocytosis by warming up, ensuring synchronous phagosome maturation between wt and ΔABC cells, despite their phagocytic defect. Five and fifteen min after uptake, surface proteins were present in phagosomes of both cell lines as a consequence of membrane internalization during phagocytosis (**Fig. 5A**). In wt cells, biotinylated proteins, as well as SibA and LmpB, were rapidly retrieved from phagosomes, reaching 50% after 30 minutes (**Figs. 5A-B**). On the other hand, plasma membrane proteins lingered in phagosomes as long as 180 minutes in ΔABC cells, suggesting a defect in retrieval of these proteins.

**Figure 5.**
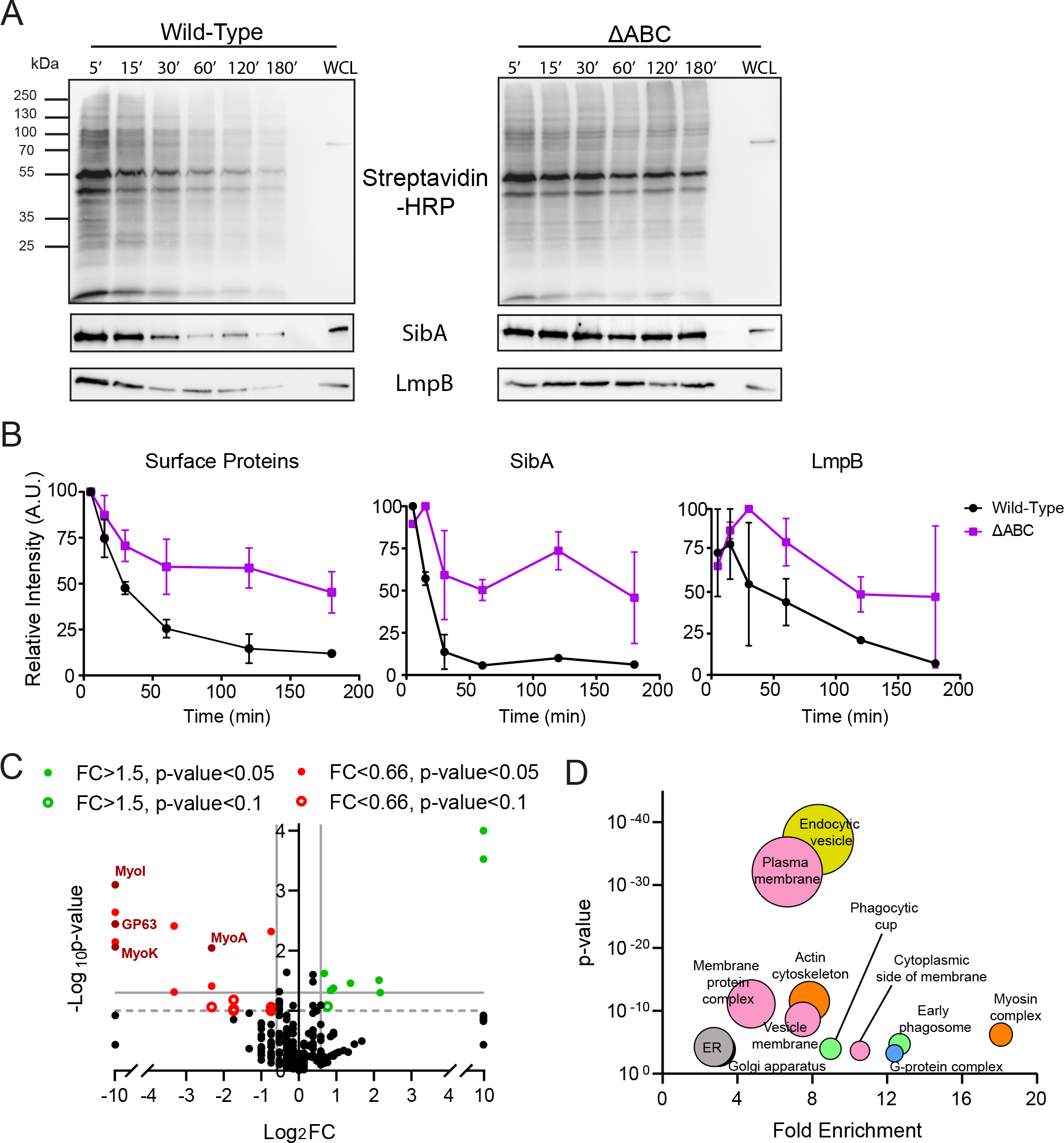
Absence of vacuolins affects plasma membrane recycling. **A-B**. Isolation of bead-containing phagosomes at different times of maturation. The cell surface of wt and ΔABC cells was biotinylated, and cells were allowed to phagocytose a large excess of latex beads for 5 or 15 minutes of pulse. After washing extracellular beads, cells were chased for the indicated time, homogenized, and phagosomes isolated by flotation on sucrose gradients. Phagosomes and the whole cell lysate (WCL) were analyzed by western blotting with Streptavidin-HRP (top) to reveal surface proteins or specific antibodies against two plasma membrane proteins (bottom). Representative blots of 2 independent experiments. **B.** Quantification of A. Relative intensity of the whole lane (Surface Proteins) or specific bands were quantified for each time point by ImageJ, and normalized to the highest value (mean ±s.e.m., N=2). **C-D.** Some specific plasma-membrane associated proteins are less abundant at the surface of ΔABC cells. The cell surface of wt and ΔABC cells was biotinylated, cells lysed and surface proteins pulled-down by streptavidin beads. Proteins were then identified by label-free mass spectrometry, and spectral counts compared between wt and mutant cells (N=3). **C.** Volcano plot showing the fold change and p-value of 335 proteins identified. Significant hits were color-coded as indicated. Note that proteins that were completely absent from wt or ΔABC cells were arbitrarily attributed the fold change of 1000 or 0.001 respectively. **D.** Bubble chart of enriched Cellular Component GO terms. GO terms were manually curated to exclude redundant terms, using the DAVID software. Terms belonging to similar clusters or components are similarly color-coded.

To determine which plasma membrane proteins may be mistrafficked in absence of vacuolins, we used biotinylation and streptavidin pull-down to identify the surface proteome of wt and ΔABC cells (**Figs. 5C-D and Sup. Tables 3-4**). Mass spectrometry analyses first identified 700 proteins. Manual curation was used to exclude nuclear, mitochondrial or ribosomal proteins, revealing 336 plasma membrane and strongly associated cortical proteins (**Fig. 5C and Sup. Table 4**). Of these, 130 proteins contain transmembrane domains, or are known to be glycosylated or lipidated. In addition, 65 proteins belonged to the “plasma membrane” cellular component GO term (**Fig. 5D**). Moreover, proteins belonging to GO terms including membrane proteins, cytoskeleton or the endocytic maturation were highly enriched in our surface proteome. This suggests that we collected not only proteins inserted into the plasma membrane, but also candidates that associate and/or interact strongly with membrane proteins. Twelve proteins were significantly under-represented in the surface proteome of ΔABC cells (**Fig. 5C and Sup. Table 3**). Of these, notably, three myosins were identified: MyoA, MyoI and MyoK. In addition, several uncharacterized proteins with transmembrane or EGF-like domains were less abundant at the surface of ΔABC cells. These results lead us to conclude that *D. discoideum* vacuolins, like mammalian flotillins, are involved in plasma membrane protein recycling, thus affecting phagocytic uptake.

### Vacuolins are involved in lysosome biogenesis but not bacterial killing

As vacuolins are PL markers, we wondered whether their depletion may affect phagosomal maturation and PL biogenesis. Acidification and proteolysis are two hallmarks of phagosome maturation, and we tested whether these functions may be impaired in vacuolin KO mutants. Wt and ΔABC cells were incubated with silica beads coupled to a pH-sensitive fluorophore (FITC) to measure phagosomal acidification over time (Sattler et al., 2013). To ensure that only efficiently phagocytosed beads were considered, cells were tracked by time-lapse microscopy and the fluorescence of individual intracellular beads was measured (**Fig. 6A**). Upon uptake, the phagosomes of both wt and ΔABC cells were rapidly acidified. In wt cells, reneutralization, which is achieved by removal of the v-ATPase, started around 45 minutes, and phagosomes reached a neutral pH about 90 min post uptake (**Fig. 6A**). Conversely, in ΔABC cells, the reneutralization phase started earlier and a neutral pH was reached as rapidly as 40 min after uptake. To monitor phagosomal proteolysis, a similar assay was used, using silica beads coupled to the self-quenched DQGreen-BSA. Proteolytic cleavage of BSA releases DQGreen molecules into the phagosomal lumen, thus increasing its fluorescence over time (Sattler et al., 2013). In wt and ΔABC cells, proteolysis started right after bead uptake, reaching a plateau after about 10 min (**Fig. 6B**). Interestingly, we observed a slight, although not striking, earlier onset of proteolysis in ΔABC cells.

**Figure 6.**
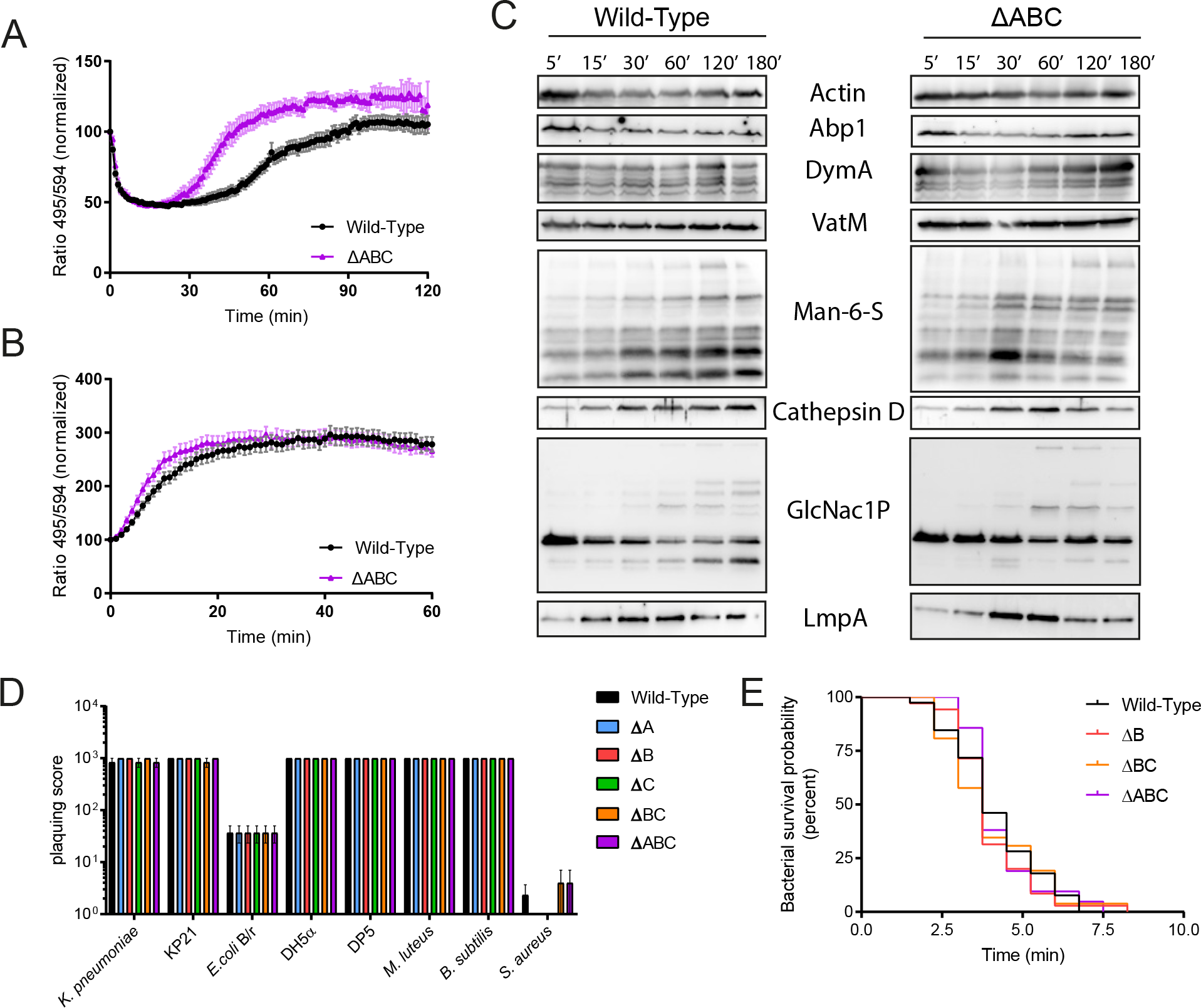
Absence of vacuolins impairs lysosome biogenesis but not bacteria killing. **A.** The reneutralization phase occurs earlier in ΔABC cells. Wt and ΔABC cells were spinoculated with silica beads labeled with the fluorophores CF594 (insensitive to pH) and FITC (sensitive to pH) and imaged for two hours by time-lapse microscopy. Beads were tracked manually using the Imaris software, and the ratio of both fluorophores normalized to time point 0 (mean ±s.e.m., n=35 beads for wt, n=19 beads for ΔABC cells, N=4). **B.** Proteolysis occurs slightly earlier in ΔABC cells. Same as in A, using beads labeled with the fluorophores CF594 and DQGreen-BSA (fluoresces upon proteolysis, mean ±s.e.m., n=31 beads for wt, n=28 beads for ΔABC cells, N=3). **C.** Lysosomal and postlysosomal markers are recruited earlier on phagosomes in ΔABC cells. Phagosomes were isolated as in **Fig. 5A** and analyzed by western blotting with the indicated antibodies. Representative blots of 2 independent experiments. **D.** Vacuolin KO mutants are able to grow on a variety of bacterial species. Different dilutions of wt or mutant cells were deposited on the indicated bacterial lawns. The plaquing score was determined using a logarithmic scale. Examples of plaques are shown in **Fig. S4** (mean ±s.e.m., N=4). **E.** Absence of vacuolins does not affect bacteria killing. Wt or mutant cells were plated in the presence of GFP-expressing *K. pneumoniae* and imaged by time-lapse microscopy at 30 sec intervals. The time from ingestion until disappearance of the GFP signal, and thus killing, was counted and a survival probability was calculated. Kaplan-Meier survival curves are shown (n≥30 events, N=2, median killing time: 3.75 min for both strains).

To better investigate a possible role of vacuolins in lysosome biogenesis, we isolated latex bead-containing phagosomes at different maturation stages of wt and ΔABC cells, as for **Fig. 5**. Then, the temporal profiles of different maturation markers were analyzed by western blotting (**Fig. 6C**). Actin and Abp1, which are associated with the phagosome right after closure, rapidly dissociated from the phagosome 15 min after uptake in wt cells. In ΔABC cells, this dynamic was similar, although we observed slightly more cytoskeleton and its associated proteins at later time points (**Fig. 6C and S4A**). The transmembrane v-ATPase subunit, VatM, had similar recruitment dynamics between wt and mutant cells. In *D. discoideum*, there are two different types of lysosomal enzymes, distinguished by their sugar modifications (Freeze et al., 1990; Souza et al., 1997). Enzymes bearing the *N*-acetylglucosamine-1-phosphate (GlcNac1P) modification are first delivered to the phagosome, followed later on by those carrying the Mannose-6-Sulfate (Man-6-S, Common Antigen-1) modification (Souza et al., 1997). Cathepsin D, among others, belongs to the latter group (Journet et al., 1999). We observed that in ΔABC cells, Man-6-S-modified enzymes appeared earlier than in wt cells, with a peak reached at 30-60 min instead of 120 min (**Fig. 6C**). Moreover, certain GlcNac1P enzymes were slightly under-represented in ΔABC phagosomes. LmpA, a PL glycoprotein involved in lysosomal biogenesis (Sattler et al., 2018), was also recruited to compartments earlier in cells lacking vacuolins (**Fig. 6C**). These results suggest that absence of vacuolins may impair delivery and/or retrieval of lysosomal proteins and thus impact lysosomal functions.

To test whether the differences in recruitment dynamics of lysosomal proteins in vacuolin KO mutants may affect digestion and killing of bacteria, we monitored growth of wt and vacuolin mutant cells on different Gram-negative and Gram-positive bacteria. For this, cells were plated in increasing dilutions on bacterial lawns and the formation of phagocytic plaques was monitored over time (**Fig. 6D and S4B**). The large excess of bacteria in the lawn bypasses the uptake defects of vacuolin KO mutants. To our surprise, vacuolin KO mutants were able to grow, digest and kill every bacterial species tested as well as wt cells. To measure intracellular bacterial killing more precisely, we monitored by time-lapse microscopy killing of GFP-expressing *K. pneumoniae*, as previously described (Leiba et al., 2017). Disappearance of the GFP signal was used as a proxy for killing (**Fig. 6E**). On average, *K. pneumoniae* was killed in 3.75 min in wt cells as well as in all vacuolin KO mutants tested.

To conclude, these results indicate that vacuolin KO mutants may have a shorter phagosomal acidic phase, as well as faster delivery or retrieval of certain lysosomal enzymes, with no apparent impact on bacterial killing and digestion.

### Absence of vacuolins impacts early recruitment of WASH and expression of *myoI*

In *D. discoideum*, the conserved WASH complex is involved in retrieval of the v-ATPase and lysosomal enzymes from lysosomal compartments to allow PL maturation, as well as in plasma membrane proteins recycling from phagosomes and macropinosomes (Carnell et al., 2011; King et al., 2013; Buckley et al., 2016). In light of the phenotypes of vacuolin depletion described here, we hypothesized that the WASH complex may be affected. To test this hypothesis, we transfected wt and ΔABC cells with GFP-WshA and analyzed its early recruitment dynamics on BCPs by time-lapse microscopy (**Fig. 7A-B**). WASH was recruited as early as 80 seconds after phagosomal closure, in both wt and KO cells, as previously published (Buckley et al., 2016). However, while in wt cells WASH was retrieved from the majority of BCPs after about 3 min, in ΔABC cells the complex stayed on the phagosome at least one more minute (**Fig. 7B**).

**Figure 7.**
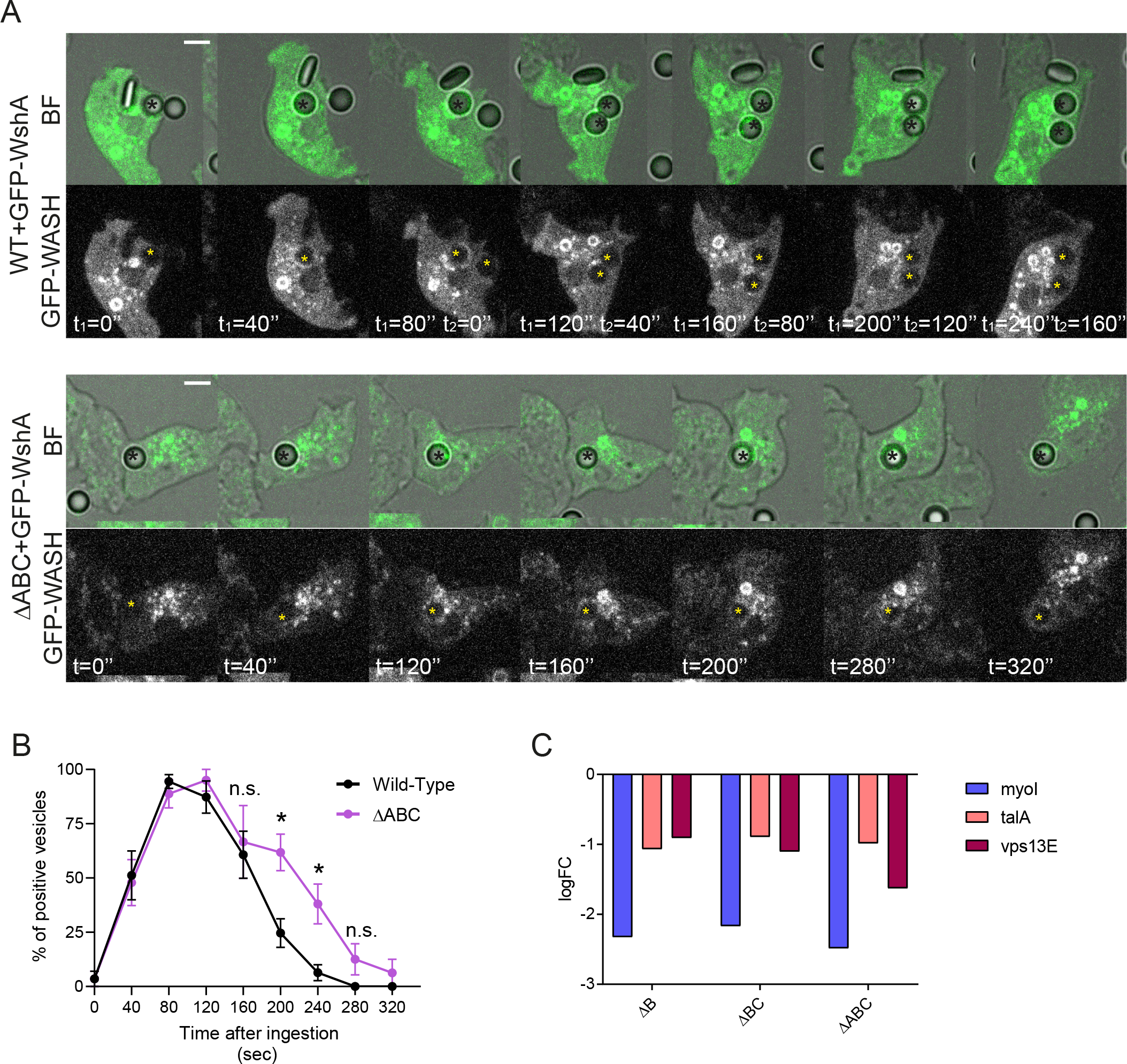
Absence of vacuolins impacts both the dynamics of WASH recruitment and the expression of three genes involved in membrane trafficking. **A.** Absence of vacuolins affects the first wave of WASH recruitment. Wt or ΔABC cells overexpressing GFP-WASH were spinoculated with latex beads and imaged by time-lapse microscopy. Representative Brightfield (BF) and GFP-WASH images are shown for each strain. Time from uptake is indicated, asterisks labels the phagocytosed bead, scale bar, 5 μm. **B.** Quantification of A. Percentage of WASH-positive vesicles was quantified at the indicated time points (N=4) **C.** *myoI*, *talA* and *vps13E* transcripts are downregulated in vacuolin KO mutants. RNA-sequencing was performed in vacuolin KO mutants, RNA levels were compared with wt cells and the log_2_(Fold Change) was plotted (N=3, p≤0.05).

As another approach to determine the mechanism by which vacuolins may affect plasma membrane recycling and lysosome biogenesis, we used a transcriptomic approach. The RNA of single, double and triple vacuolin KO mutants was collected, sequenced and compared with wt cells (**Fig. 7C**). We found that absence of one or several vacuolins affected transcription of *myoI.* Strikingly, MyoI was also completely absent from the surface proteome of ΔABC cells (**Fig. 5C**). Interestingly, KO of *myoI* induces a severe phagocytosis defect similar to the one described here (Titus, 1999). Talin A, an actin binding protein, was shown to interact with MyoI and be downregulated upon MyoI depletion (Niewöhner et al., 1997a; Gebbie et al., 2004; Tuxworth et al., 2005). Accordingly, we find that in vacuolin KO strains, transcription of *talA* is also affected, probably as a consequence of *myoI* downregulation (**Fig. 7C**). Moreover, the transcription of *vps13E* (vacuolar protein sorting 13 E) is also highly affected in all vacuolin KO strains tested when compared with wt cells (**Fig. 7C**). These results suggest that vacuolins may act in concert with the WASH complex in plasma membrane protein recycling. Furthermore, our RNAseq suggest that absence of vacuolins affects transcription of membrane trafficking proteins, which may result in the defects we report here.

## DISCUSSION

Metazoan flotillins are lipid rafts proteins that function as signaling platforms at the plasma membrane and in membrane trafficking of an increasing number of cargoes in the endocytic pathway (Otto and Nichols, 2011; Meister and Tikkanen, 2014). Vacuolins have been proposed to act as the flotillin homologues in the amoeba *D. discoideum* based on protein structure similarities and sequence conservation of the PHB domain (**Fig. S1F**, (Rauchenberger et al., 1997; Jenne et al., 1998)). In this study, we further confirm that all three vacuolins share the same biochemical characteristics as mammalian flotillins, in that they partially associate with alkali- and detergent-resistant membranes (**Fig. 2**). Flotillins insert into the inner leaflet of membranes through post-translational palmitoylations occurring in the PHB domain, which are a prerequisite for membrane association (Morrow et al., 2002; Neumann-Giesen et al., 2004). The PHB domain of vacuolins was also shown to be required for membrane association, as evidenced by deletion mutants lacking this domain (Wienke et al., 2006). It is important to note that the PHB domain of each vacuolin possesses two cysteines (**Fig. S1F**), and we speculate that these may also be palmitoylated to allow membrane association. In addition, like flotillins, vacuolins have been shown to oligomerize via protein-protein interactions of the C-terminal coiled-coil regions and proposed to form trimers (Wienke et al., 2006; Solis et al., 2007). On the other hand, vacuolins and flotillins also share some differences. Namely, we find that vacuolins only partially associate with DRMs (**Fig. 2C**). Interestingly, the membrane insertion topology of flotillins and vacuolins may differ, as the PHB domain of flotillins is at the N-terminus of the protein, whereas in vacuolins the PHB domain is more central (**Fig. S1F**). Moreover, although flotillins are found across several compartments, from the plasma membrane to endosomal vesicles and the Golgi apparatus (Babuke and Tikkanen, 2007), we mainly detected vacuolins on endocytic compartments and the contractile vacuole, and only transiently at the plasma membrane (**Fig. 1**). How vacuolins are targeted to their destination compartment remains currently unknown.

Thanks to the generation of vacuolin KIs, we have demonstrated that vacuolins not only accumulate on PLs, but are recruited to phagosomes very quickly after phagosomal closure (**Fig. 1D**). Since no vacuolin signal was detected on BCPs during the formation of the phagosome, we suggest that vacuolins are probably delivered to the phagosomes via fusion with vacuolin-positive vesicles.

We find that deletion of one or more vacuolin severely impairs uptake, independently on the nature of the particle (**Fig. 3**). In addition, we provide evidence that absence of vacuolins affects particle recognition, and perhaps also adhesion, (**Fig. 4D-E**), but does not significantly affect actin-mediated motility (**Fig. 4C**) or formation and closure of the phagocytic cup (**Fig. 4A-B**). This severe uptake defect is reminiscent of the phenotype of the KO strain of the unconventional myosin MyoVIIa/MyoI (Titus, 1999). MyoI depletion does not alter formation or closure of the phagocytic cup, but affects adhesion to particles and substrate, thereby impairing motility, migration and uptake (Tuxworth et al., 2001; Gebbie et al., 2004; Tuxworth et al., 2005). The interacting partner of MyoI, the talin TalA, is also necessary for efficient substrate and particle adhesion and thus exhibits a similar uptake defect (Niewöhner et al., 1997b; Gebbie et al., 2004; Tuxworth et al., 2005). Interestingly, we find that in vacuolin KO cells, both genes are downregulated (**Fig. 7C**) and consequently, MyoI was severely depleted from the plasma membrane-associated proteome in these cells (**Fig. 5C**). However, it is important to note that vacuolin KO cells do not display substrate adhesion, motility or cytokinesis defects as severe as *myoI* and *talA* KO cells (Niewöhner et al., 1997b; Gebbie et al., 2004). This indicates that the uptake phenotype we observe is more specific than the *myoI* or *talA* KOs. For example, vacuolins may be required for the proper function of these proteins only in the context of adhesion and/or in the particle recognition step of uptake. How depletion of vacuolins alters expression of these actin-interacting proteins remains to be elucidated.

Flotillins have recently been involved in the recycling of plasma membrane proteins through the Rab11-positive ERC in several cell types (Solis et al., 2013; Bodrikov et al., 2017b). We show here that KO of all three vacuolins leads to a delayed retrieval of surface proteins during phagocytosis (**Fig. 5**), which could also partly explain the uptake and adhesion defects found in these cells (**Fig. 3**). In *D. discoideum*, the WASH complex is recruited right after phagosome and macropinosome closure (Buckley et al., 2016). This early wave of WASH recruitment was suggested to be important for plasma membrane protein retrieval as WASH KO impaired recycling of surface proteins, leading to an inefficient uptake of yeasts (Buckley et al., 2016). The phenotype in surface protein recycling observed in vacuolin KO cells is very similar to the one reported for WASH KOs (Buckley et al., 2016). In addition, upon depletion of vacuolins, we find that, although the delivery of WASH at the early phagosome is not affected, the complex resides longer on the BCP (**Fig. 7A-B**). This could indicate that the WASH complex is retrieved later in absence of vacuolins: for example, vacuolins may be involved in the recycling of the complex itself. Alternatively, the function of WASH in promoting actin polymerization may be impaired in absence of vacuolins and therefore the complex lingers on BCPs.

Analysis of the surface proteome of wt and ΔABC cells highlighted three myosins that were under-represented or absent from the surface of vacuolin KOs (**Fig. 5C**). Myosins are actin-binding motors that play crucial roles in different steps of phagocytosis, such as formation of lamellipods, closure of the cup and trafficking of vesicles to the phagosome in mammals (Araki, 2006). In *D. discoideum*, MyoB was shown to participate in plasma membrane recycling (Neuhaus and Soldati, 2000). MyoK, one of the hits we identified, is found at the phagocytic cup and is involved in cup closure as well as in trafficking of ER and lysosomal proteins to the phagosome (Dieckmann et al., 2010, 2012). These data suggest that, when vacuolins are depleted, certain myosins may be mistrafficked and therefore impact uptake and/or correct phagosomal maturation.

Interestingly, upon vacuolin overexpression, cells were also impaired in uptake (**Fig. 3D**). Flotillins assemble on specific microdomains, however when they are overexpressed, newly formed flotillin-microdomains can be observed at the plasma membrane (Frick et al., 2007). In our case, we observed an increased number of vacuolin- and p80-positive compartments, suggesting that overexpression of vacuolins may affect formation and/or fusion of PLs (**Fig. S1C**). As vacuolins assemble in oligomers, we suggest that overexpression of one of them may cause an imbalance in the stoichiometry of the oligomers, thus affecting their function.

Finally, we show here that vacuolins are involved in postlysosome biogenesis, in that, in their absence, the reneutralization phase occurs earlier and lysosomal enzymes and PL proteins are found earlier in phagosomes (**Fig. 6A-C**). It is possible that vacuolins are involved in the delivery and/or retrieval of lysosomal and PL proteins from phagosomes, by creating specific sorting areas on the compartment. Whether vacuolin absence affects the WASH-dependent recycling of the v-ATPase and lysosmal enzymes has not been tested yet. Despite this faster maturation, phagosomes of vacuolin KO mutants are still bactericidal and as proteolytic as in wt cells. This confirms once again, that acidification is not the only factor required for killing bacteria, which also requires proteolysis, oxidation and/or toxic metals (Cosson and Lima, 2014; Barisch et al., 2018).

Surprisingly, the phenotypes described here do not reproduce what had been found with the previously described *vacA-* and *vacB-* mutants (Jenne et al., 1998). It is possible that these phenotypes are specifically found in the Ax2(Ka) *D. discoideum* background, however we have shown that the *vacB-* mutant had deletions of several genes (**Fig. S3**), which may have altered the original description of the effect of *vacB* KO. To conclude, here we report that vacuolins behave and partially function like the flotillin homologues in *D. discoideum*, indicating that the role of these proteins in plasma membrane recycling may be conserved throughout evolution.

## MATERIALS AND METHODS

### *D. discoideum* strains, culture and plasmids

*D. discoideum* strains and plasmids are described in Supplementary Table 1. Cells were axenically grown at 22°C in HL5c medium (Formedium) supplemented with 100 U/mL of penicillin and 100 μg/mL of streptomycin (Invitrogen).

To generate GFP knock-ins, 5’ and 3’ homologous regions of each vacuolins were cloned using the primers described in table S2. The arms were cloned using BglII/SpeI for the left arm (LA) and SalI/SacI for the right arm (RA) into the plasmid pPI183, which allows targeted in-frame integration of GFP at the C-terminus of the gene of interest (Paschke et al., 2018). Integration was confirmed using flanking primers described in Table S2 by PCR and by western blot. In addition, the level of expression of KI genes was compared with that of wt cells by western blot and qRT-PCR to ensure comparable levels of expression. To generate single vacuolins knock-out strains, 5’ and 3’ homologous regions of each vacuolins were cloned using the primers described in Table S2 into the pKOSG-IBA-dicty1 plasmid following manufacturer’s instructions (Stargate). Knock-out was confirmed using flanking primers described in table S2 by PCR and qRT-PCR. The BsR cassette was then floxed from single mutants using the Cre recombinase. To generate double simultaneous ΔBC strains, the left arm of VacC and the right arm of VacB were cloned into pKOSG and clones were verified as before. To generate the triple ΔABC strain, the plasmid pJET-VacA-Hyg was transfected into ΔBC strain and verified as before. Plasmids and primers used to generate GFP-tagged vacuolins are listed in table S1 and S2. Plasmids were transfected into *D*. *discoideum* by electroporation and selected with the relevant antibiotic. Hygromycin was used at a concentration of 15 μg/mL, blasticidin and G418 were used at a concentration of 5 μg/mL.

### Bacterial strains and culture

The bacterial strains used in this study are listed in Table S1. Bacteria were cultured in LB medium at 37°C in shaking as previously described (Froquet et al., 2009). Mycobacteria were grown in Middlebrook 7H9 (Difco) supplemented with 10% OADC (Becton Dickinson), 0.2% glycerol and 0.05% Tween 80 (Sigma Aldrich) at 32°C in shaking culture at 150 r.p.m in the presence of 5 mm glass beads to prevent aggregation.

### Antibodies, reagents, western blotting and immunofluorescence

Recombinant nanobodies with the Fc portion of rabbit IgG that recognize specifically VacA or VacB were obtained via the Geneva Antibody Facility (University of Geneva). Cross-reaction and specificity were tested by immunofluorescence on knock-out strains. Note that these antibodies only work in immunofluorescence and that unfortunately, we were not able to produce VacC-specific antibodies. **Fig. 2** illustrates that the 221-1-1 pan-vacuolin antibody is not able to discriminate between the different vacuolins. The other following antibodies were used: pan-vacuolin (221-1-1, Dr. M. Maniak (Jenne et al., 1998)), VatA (Dr. M. Maniak (Jenne et al., 1998)), VatM (N2, Dr. R. Allen (Fok et al., 1993)), actin (Dr. G. Gerisch (Westphal et al., 1997)), Abp1 (Dieckmann et al., 2010), LmpA and LmpB (Dr. M. Schleicher (Janssen et al., 2001)), mitochondrial porin (Dr. G. Gerisch), p80 (purchased from the Geneva Antibody Facility), SibA (Dr. P. Cosson (Cornillon et al., 2006)), DymA (Dr. D. J. Manstein (Wienke et al., 1999)), Common Antigen-1 (Dr. M. Maniak), GlcNac1P (AD7.5, Dr Hudson Freeze (Souza et al., 1997)), cathepsin D (Dr. J. Garin (Journet et al., 1999)) and GFP (pAb from MBL Intl., mAb from Abmart). Goat anti-mouse or anti-rabbit IgG coupled to AlexaFluor488 or AlexaFluor594 (Invitrogen) or to HRP (Brunschwig) were used as secondary antibodies.

The lipophilic membrane dye FM4-64 (*N*-(3-Triethylammoniumpropyl)-4-(6-(4-(Diethylamino) Phenyl) Hexatrienyl) Pyridinium Dibromide, Invitrogen) was used at a concentration of 1 μg/mL.

After SDS-PAGE separation and transfer onto nitrocellulose membranes (Protran, Schleicher & Schuell), immunodetection was performed as previously described (Schwarz et al., 2000) but with ECL Prime Blocking Reagent (Amersham Biosciences) instead of non-fat dry milk. Detection was performed with ECL Plus (Amersham Biosciences) using a Fusion Fx device (Vilber Lourmat). Quantification of band intensity was performed with ImageJ.

For immunofluorescence, *D. discoideum* cells were fixed with ultra-cold methanol (MeOH) and immunostained as previously described (Hagedorn et al., 2006). Images were recorded with a Leica SP8 confocal microscope using a 63×1.4 NA or a 100×1.4 NA oil immersion objectives.

### Live imaging

Cells were plated on a μ-dish (iBIDI) in filtered HL5c, to decrease autofluorescence. After adherence, either 0.6 μm sections or time-lapse movies were taken with a spinning disc confocal system (Intelligent Imaging Innovations) mounted on an inverted microscope (Leica DMIRE2; Leica) using the 63 × or the 100 × 1.4 NA oil objective. Images were processed with ImageJ. Quantifications were performed manually. A 1 mm-thin agarose sheet was overlayed onto cells before time-lapse microscopy to allow better imaging of the phagosome.

### Motility assay by high-content microscopy

*D. discoideum* cells were plated overnight at a density of 2×10^5^ cells/mL in filtered HL5c. The next day, about 2×10^4^ cells diluted in Sorensen-Sorbitol (14.7 mM KH_2_PO_4_, 2.5 mM NaHPO_4_, pH 6.2, 120 mM sorbitol) were deposited in 96-well plates (Perkin Elmer) and left to attach for 30 min at room temperature. Medium was then aspirated and replaced by Sorensen-Sorbitol alone or with 2 mM Folate, and cells were incubated for 20 min. Images were taken every 15 seconds for 30 minutes with a 10x objective with the ImageXpress Micro XL high-content microscope (Molecular Devices). Cells were tracked with the MetaXpress software (Molecular Devices) and average speed was calculated with MS Office Excel.

### Particle uptake

Assessment of fluorescent particles uptake was performed as previously described (Sattler et al., 2013). Data was acquired using a FACSCalibur Flow cytometer (BD Biosciences) or FACS Gallios (Beckman Coulter) and analyzed with the FlowJo software (TreeStar) or Kaluza (Beckman Coulter).

### Acidification and proteolysis

The kinetics of acidification and proteolytic activity inside phagosomes were monitored as previously described (Sattler et al., 2013). Phagosomal pH was monitored using the pH-sensitive fluorophore FITC (Molecular Probes) coupled to 3 µm silica particles (Kisker Biotech). Beads were also coated with the pH-insensitive CF594 (Biotium) as a reference dye. Cells were plated in HL5c filtered on 35-mm iBIDI dishes. 30 minutes before imaging, HL5c was replaced by LoFlo (Formedium). Beads were added at a 1:2 ratio and a 1 mm-thin agar sheet was overlayed on top of the cells. Movies were recorded with a Leica AF6000LX widefield microscope using a 40x objective. Images were taken every 1 min for 4 hours. Tracking of bead-containing phagosomes and calculations of their emitted fluorescence ratios were done with the Imaris software (Bitplane). Proteolysis was measured using beads coupled to the reference stain CF594 (Biotium) and the reporter DQ Green-BSA (Molecular Probes) at a self-quenching concentration. Upon proteolysis of BSA, DQ Green is released and dequenched, which causes an increase in fluorescence. Movies were recorded as above.

### Phagocytic plaque assay

The ability of *D. discoideum* to form plaques on a lawn of bacteria was monitored as previously described (Froquet et al., 2009). Briefly, 50 µl of an overnight bacterial culture was plated on SM-agar (10 g peptone (Oxoid), 1 g yeast extract (Difco), 2.2 g KH_2_PO_4_, 1 g K_2_HPO_4_, 1 g MgSO_4_ × 7H_2_O) with 20% glucose in 1 L of ddH_2_O) in wells from a 24-well plate. Serial dilutions of *D. discoideum* (10, 10^2^, 10^3^ or 10^4^ cells) were plated onto the bacterial lawn, and the plates were incubated at 21°C for 4-7 days until plaque formation was visible. To quantify cell growth on bacteria, a logarithmic growth score was assigned as follows: plaque formation up to a dilution of 10 cells received a score of 1000; when cells were not able to grow at lower dilutions, they obtained the corresponding lower scores of 100, 10 and 1.

### Intracellular killing of *K. pneumoniae*

Intracellular killing of individual bacteria was measured as previously described (Leiba et al., 2017). Briefly, *K. pneumoniae*-GFP bacteria were mixed with *D. discoideum* cells at a ratio of 3:1 in Sorensen-Sorbitol, deposited on an iBIDI dish for 10 min, then imaged every 30 sec for 2-3 h with a Leica AF6000LX widefield microscope using a 40x objective with the autofocus function. Images were processed and quantified with ImageJ. Survival analysis of phagocytosed fluorescent bacteria was computed using the Kaplan–Meier estimator.

### Cytosol-membrane separation and membrane treatments

10^9^ *D. discoideum* cells were washed in Sorensen-Sorbitol and resuspended in HESES buffer (HEPES 20 mM, 250 mM Sucrose, MgCl_2_ 5 mM, ATP 5 mM) supplemented with proteases and phosphatase inhibitors (cOmplete EDTA-free and PhosSTOP, Roche). Cells were homogenized in a ball homogenizer with 10 μm clearance. The post-nuclear supernatant was diluted in HESES buffer and centrifuged at 35’000 rpm in a Sw60 Ti rotor (Beckmann) for 1 hour at 4°C. The cytosol (supernatant) and membrane (MB, pellet) fractions were recovered. The membrane fraction was further treated for 1 h on ice by addition of: HESES buffer (negative control), 1 M Urea (in HESES), 500 mM NaCl (in HESES) or 200 mM Sodium Carbonate at pH 11 (pure) in the same final volume as the recovered cytosol fraction (about 3 ml). Membranes were centrifuged at 35’000 rpm in a Sw60 Ti rotor for 1 hour at 4°C. The supernatant (SN) and pellet (P) fractions were recovered. After acetone precipitation of the SN, both SN and P fractions were resuspended in equal volumes (1.5 ml each). The protein concentration of the cytosol fraction was further quantified by Bradford and an equivalent (1:1) amount of MB, SN and P fractions were loaded for western blotting.

### Detergent-resistant membrane isolation

10^9^ *D. discoideum* cells were washed in Sorensen-Sorbitol and resuspended in 1 ml of cold Lysis Buffer (Tris-HCl pH 7.5 50 mM, NaCl 150 mM, Sucrose 50 mM, EDTA 5 mM, ATP 5 mM, DTT 1 mM) with 1% Triton X-100 and supplemented with proteases and phosphates inhibitors. The lysate was then incubated at 4°C on a rotating wheel for 30 min. After centrifugation (5 min, 13’000 rpm, table-top centrifuge, 4°C), the supernatant (Triton Soluble Fraction, TSF) was collected and the pellet (Triton Insoluble Fraction, TIF) was resuspended in 200 μl of cold Lysis buffer without Triton X-100. The TIF was mixed with 800 μl of 80% sucrose (final concentration 65% sucrose), deposited at the bottom of an ultracentrifuge tube (Beckmann polyallomer for Sw60) and overlayed with 2 ml of 50% sucrose and 1 ml of 10% sucrose. The TIF was then centrifuged at 55’000 rpm in a Sw60 Ti rotor (Beckmann) for 2 h at 4°C. The Triton Insoluble Floating Fraction (IFF) and Non-Floating Fraction (TINFF) were collected, acetone precipitated and resuspended in Laemmli Buffer. Equivalent amounts of each fractions were loaded for western blotting.

### Phagosomes purification

8 × 10^9^ *D. discoideum* cells were washed once in Sorensen-Sorbitol buffer at pH 8 at 4°C. The cell surface was then biotinylated by incubating cells for 3 min with 30 mg of EZ-Link-Sulfo-NHS-LC-Biotin (Thermo Scientific) in Sorensen-Sorbitol buffer at pH 8 on ice. After this step, latex beads were added and phagosomes were purified at different stages of maturation exactly as described in (Gotthardt et al., 2006). Equivalent amounts of proteins from each time point were loaded for western blotting.

### Surface biotinylation and mass spectrometry

2 × 10^7^ *D. discoideum* cells were washed twice in Sorensen-Sorbitol buffer at pH 6 at 4°C. The cell surface was biotinylated by incubating cells for 10 min with 1 mg of EZ-Link-Sulfo-NHS-LC-biotin (Thermo Scientific) in Sorensen-Sorbitol buffer at pH 8 on ice. Cells were pelleted and incubated 5 min on ice in PBS containing 10 mM of glycin to quench unbound biotin. Cells were carefully washed 4 times in Sorensen-Sorbitol buffer pH 6, then lysed in RIPA buffer and incubated O/N with SpeedBeads Neutravidin magnetic beads (GE Healthcare). Pulled-down proteins and beads were then washed twice in RIPA, followed by a 15 min wash in 6 M urea and 2 washes in RIPA. Beads were then washed and collected in PBS and sent for mass spectrometry analysis. Protein were digested on-beads and peptides were analysed by nanoLC-MSMS using an easynLC1000 (Thermo) coupled with a Qexactive Plus mass spectrometer (Thermo). Database search was performed with Mascot (Matrix Science) using the *Dictyostelium discoideum* reference proteome database from Uniprot. Data were analysed and validated with Scaffold (Proteome Software) with quantitation based on spectral counting and normalized spectral abundance factors (NSAF) with statistical *t*-test. GO-term enrichment analysis was performed using PANTHER.

### RNA extraction and RNAseq

One 80% confluent dish of *D. discoideum* cells was collected and centrifuged. The cell pellet was resuspended in 400 μl of Trizol (Invitrogen # AM9738) and RNA extracted using the Direct-zol RNA MiniPrep kit according to manufacturer’s instructions (Zymo Research). The quality and the quantity of RNA was confirmed with a Bioanalyzer (Agilent, RNA 6000 Nano Kit) and Qubit 2.0 fluorometer (Thermo Scientific). Libraries were constructed from 100 ng of RNA using the Ovation Universal RNA-Seq System kit (Nugen). The quality of the libraries was verified by TapeStation (Agilent, High Sensitivity D1000 ScreenTape). Samples were pooled and run in single read 50 flow cell (Illumina) and run on a Hiseq 4000 (Illumina).

## ACKNOWLEDGEMENTS

We gratefully acknowledge Dr. P. Cosson (University of Geneva) for discussions and suggestions and the staff of the Bioimaging Center for Microscopy, the FACS core facility, the Proteomics core facility and the IGE3 Genomics Platform at the Faculty of Sciences and Faculty of Medicine of the University of Geneva for their precious help. We thank J. King for sharing plasmids and D. Moreau and the ACCESS Geneva Imaging Facility of the University of Geneva for help with the high content microscope experiments. This work was supported by multiple grants from the Swiss National Science Foundation.

---

**Figure S1.**
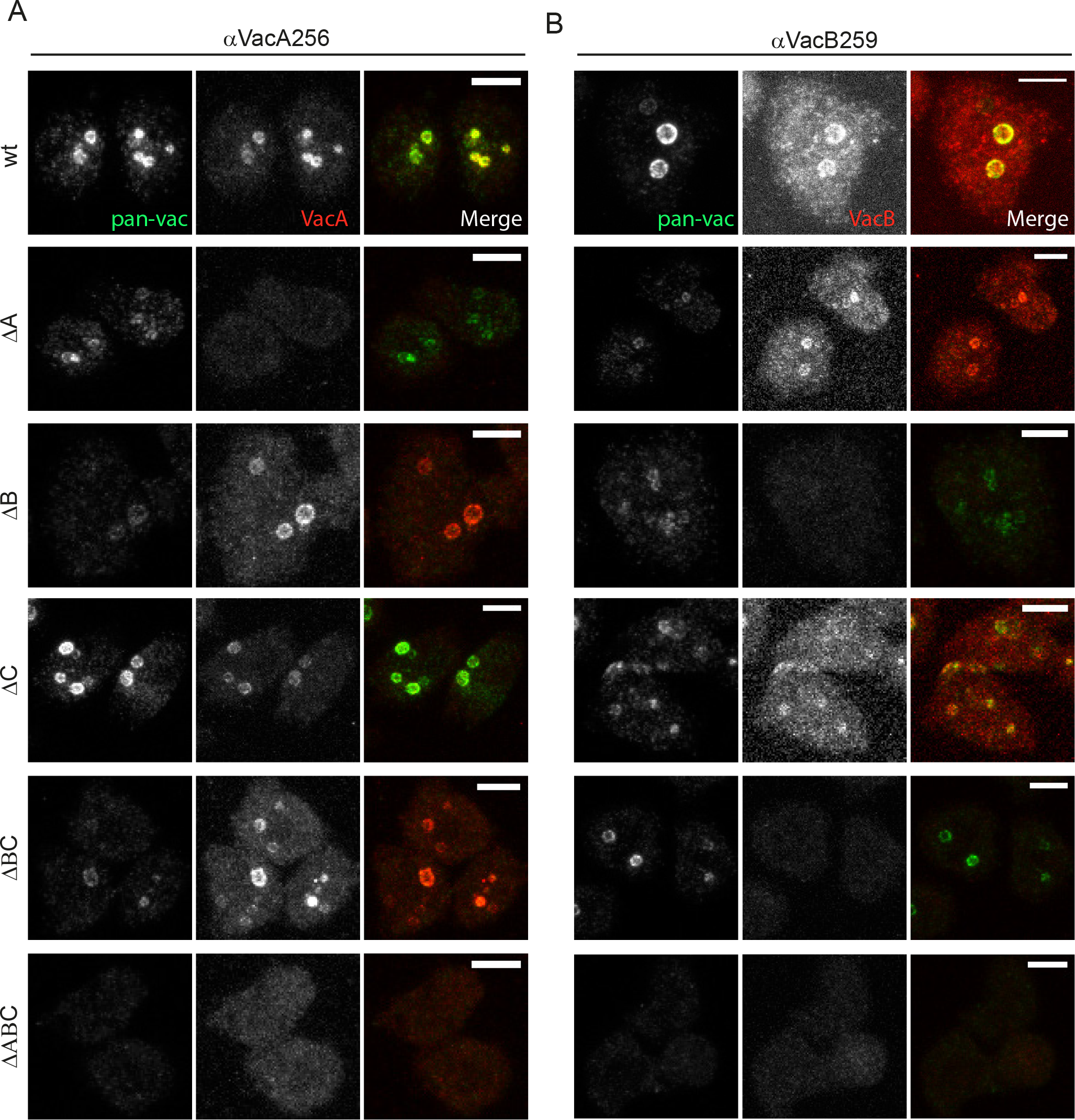
Characterization of anti-vacuolin antibodies. Wt as well as vacuolin mutant cells were fixed and immunostained with nanobodies against VacA (**A.**) and VacB (**B.**). Each nanobody is specific for its protein and does not cross-react with the other vacuolin. Scale bars, 5 μm.

**Figure S2.**
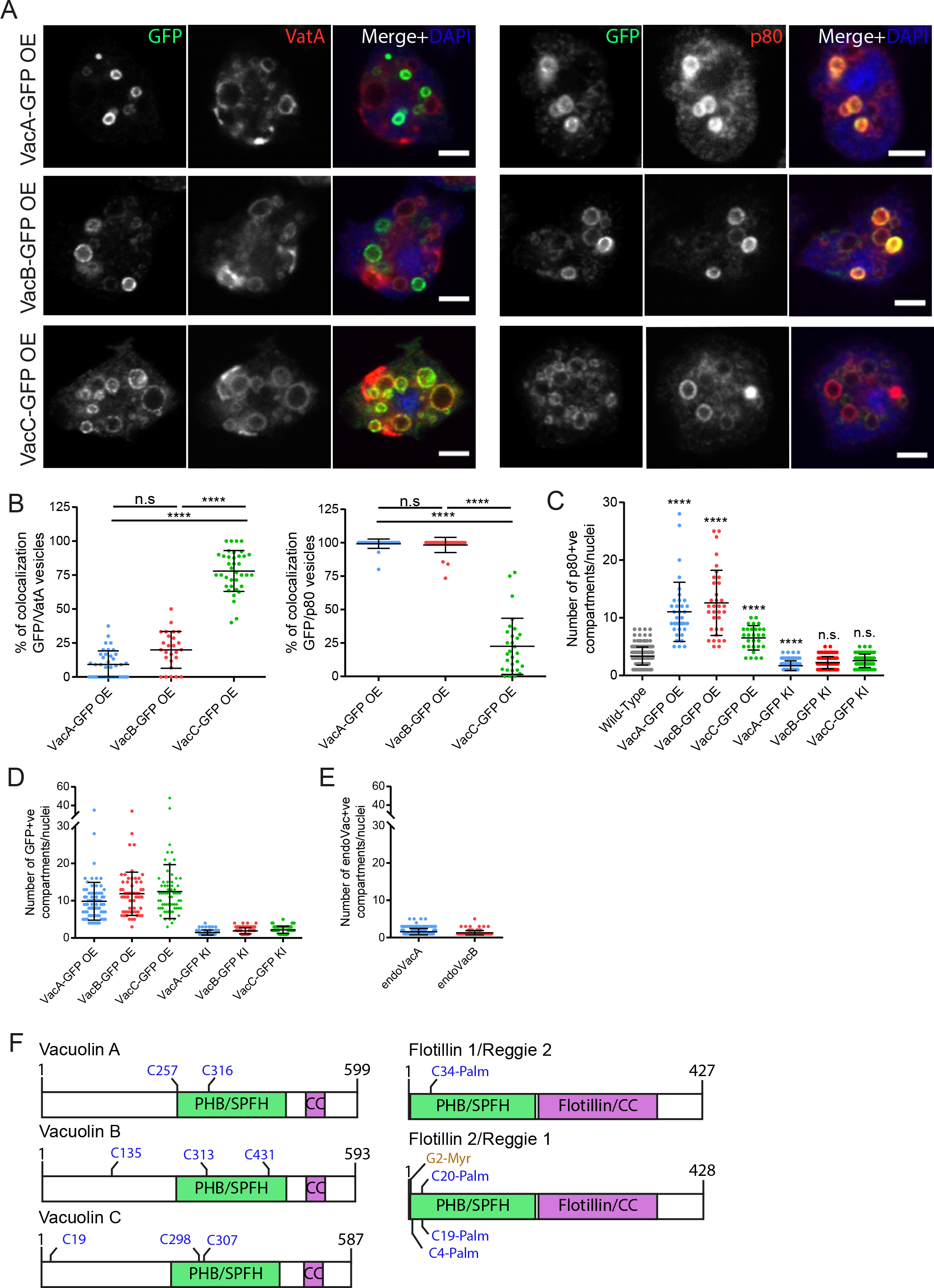
Overexpression of vacuolins affects their localization and the endosomal pathway. **A-B. Wt** cells overexpressing VacA-GFP, VacB-GFP or VacC-GFP were fixed and immunostained for GFP (green), VatA (red, left panel) or p80 (red, right panel), as well as DAPI (blue). Maximum projections are shown. Scale bars, 5 μm. **B.** Quantification of A. The percentage of colocalization was calculated for each cell, by counting the proportion of GFP positive vesicles that were VatA or p80 positive. VatA positive CV structures were not quantified. Each dot represents one cell (mean ±sd, n≥40 cells, N=1). **C.** Overexpression of vacuolins increases the number of p80 positive compartments. Quantification of the number of p80 positive compartments per cell and per nuclei. Wt, OE or KI strains were used. Each dot represents one cell (mean ±sd, n≥40 cells, N=1). **D and E.** KI of GFP does not affect the number of vacuolin positive compartments. Quantification of the number of GFP (D.) or endogenous VacA or VacB (E.) compartments per cell and per nuclei. Each dot represents one cell (mean ±sd, n≥60 cells, N=1). **G.** Scheme of the protein domains of vacuolins compared with flotillins. All the cysteines present in the sequence of each vacuolin is shown.

**Figure S3.**
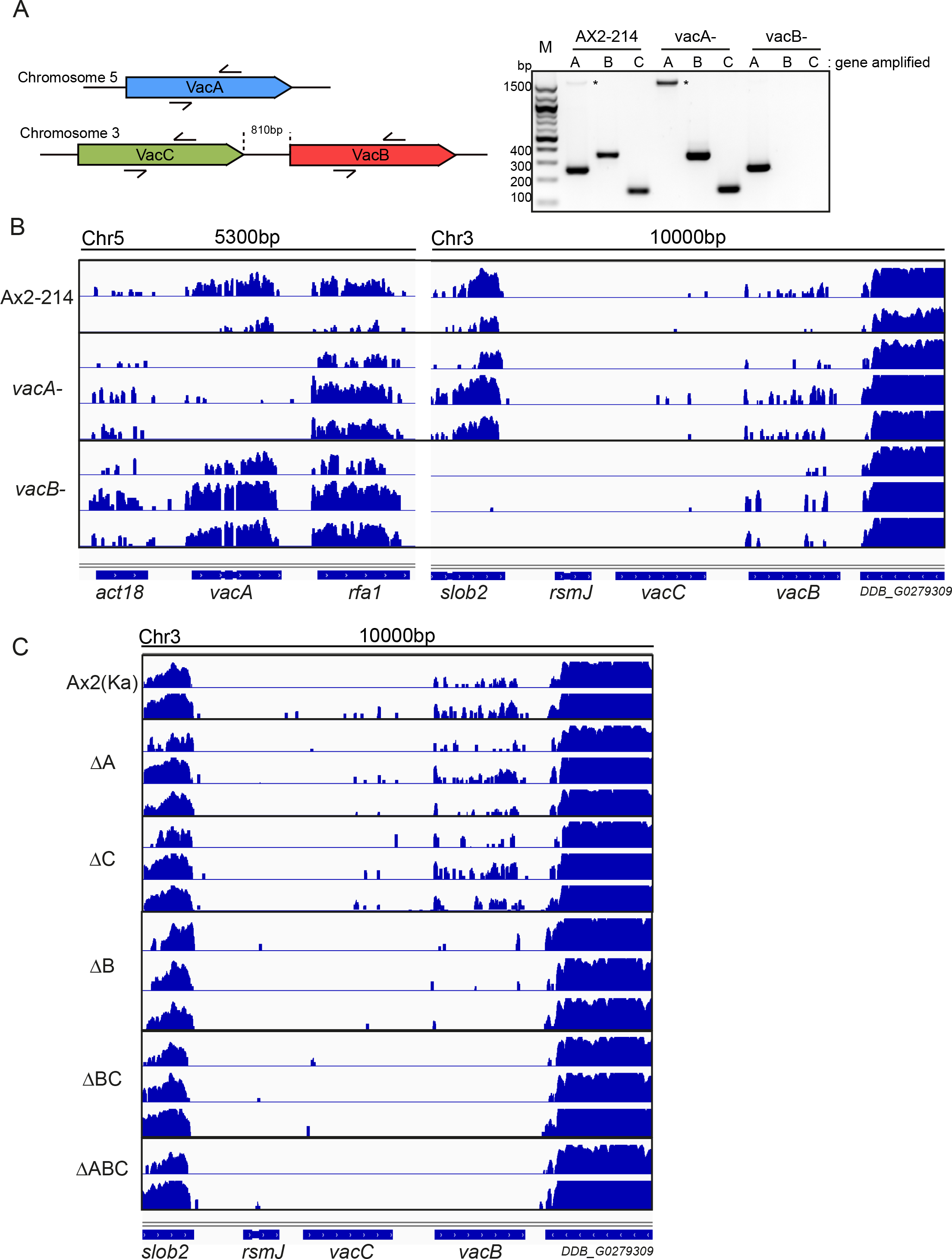
The former published vacB-mutant is not a single gene KO. **A.** Scheme of PCR primers used in PCR on genomic DNA of wt AX2-214, *vacA-* and *vacB-* cells. The *vacB-* cells lack the central portion of VacC. *, unspecific band; M, DNA marker. **B.** Read alignment obtained with the IGV software of RNA-sequencing experiments from samples isolated from AX2-214, *vacA-* and *vacB-*. Left: alignment on the *vacA* genomic region (chr 5). Right: alignment on the *vacC/vacB* genomic region (chr 3). *vacB-* cells lack reads over the *slob2* gene. **C.** Read alignment of sequenced RNA isolated from AX2(Ka) and the newly generated KO strains, obtained with the IGV software. The *slob2* gene is intact in all KOs.

**Figure S4.**
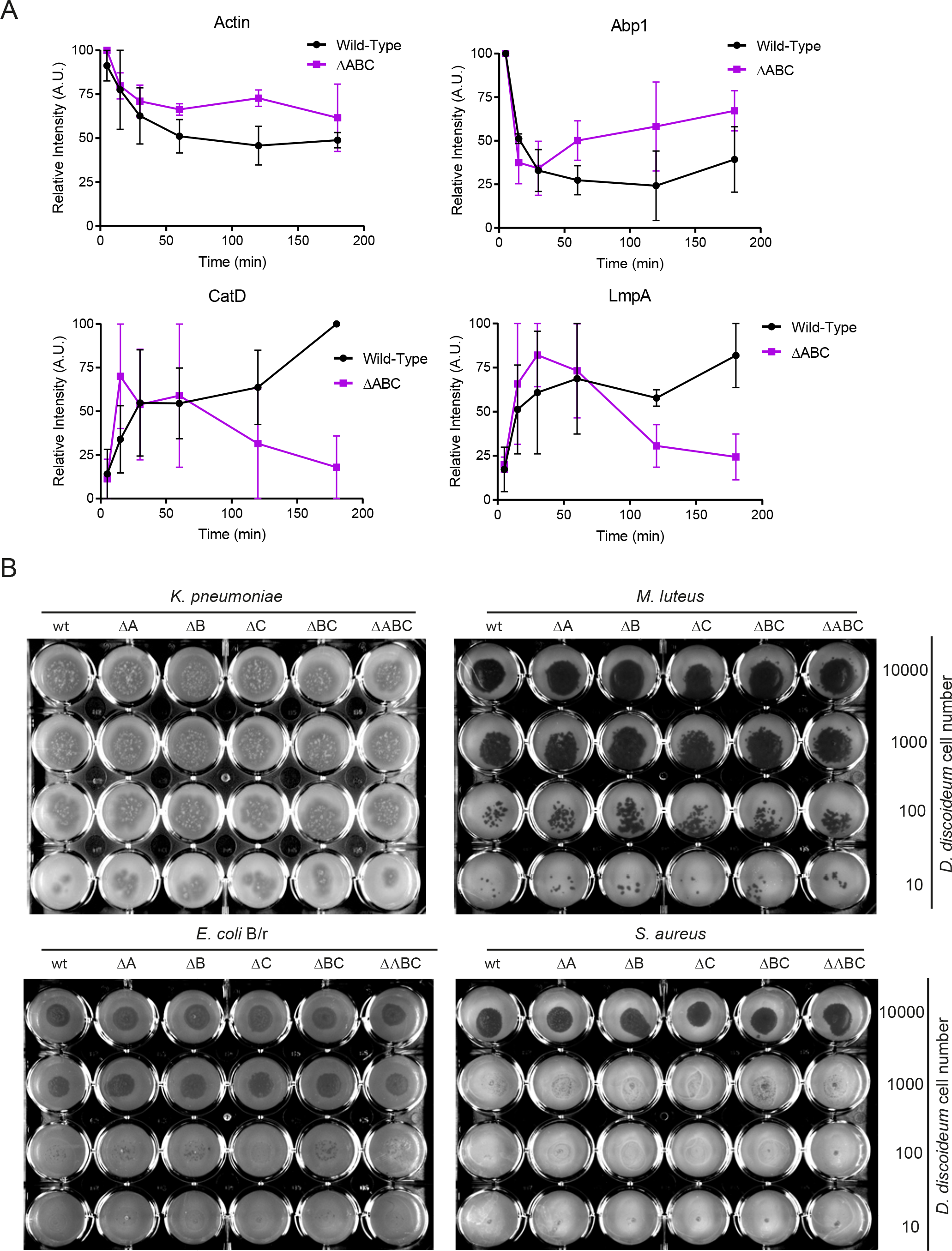
Vacuolins are involved in phagosome maturation. **A.** Quantification of Fig. 6C. Relative intensity of specific bands were quantified by ImageJ, and normalized to the highest value for each time point (mean ±s.e.m., N=2). **B.** Examples of plaque assays of Fig. 6D.

**Sup. Table 1.** List of *D. discoideum* strains used in this study.

| Strain/Plasmid | Relevant characteristics | Source/Reference |
| --- | --- | --- |
| *D. discoideum* |  |  |
| Ax2(Ka) | Wild-type |  |
| Ax2(Gerisch) | Wild-type |  |
| Ax2(Gerisch) *vacA*- knock-out strain |  | (Jenne et al., 1998) |
| Ax2(Gerisch) *vacB*- knock-out strain |  | (Jenne et al., 1998) |
| Ax2(Ka) ΔVacA knock-out strain | | This study |
| Ax2(Ka) ΔVacB knock-out strain | | This study |
| Ax2(Ka) ΔVacC knock-out strain | | This study |
| Ax2(Ka) ΔVacBΔVacC knock-out strain | | This study |
| Ax2(Ka) ΔVacAΔVacBΔVacC knock-out strain | | This study |
| Ax2(Ka) VacA-GFP knock-in strain |  | This study |
| Ax2(Ka) VacB-GFP knock-in strain |  | This study |
| Ax2(Ka) VacC-GFP knock-in strain |  | This study |
| *M. marinum* |  |  |
| M strain | Wild-type | L. Ramakrishnan (Washington University) |
| *D. discoideum* plasmids |  |  |
| pDM323 | G418<sup>r</sup>, C-terminal GFP | (Veltman et al., 2009) |
| VacA-GFP | *vacA* (DDB_G0289485) in pDM323 | This study |
| VacB-GFP | *vacB* (DDB_G0279191) in pDM323 | This study |
| VacC-GFP | *vacC* (DDB_G0279307) in pDM323 | This study |
| pPI183 | Blasticidin<sup>r</sup>, C-terminal GFP for knock-in | (Paschke et al., 2018) |
| pPI183 VacA KI | Regions of *vacA* in pPI183 | This study |
| pPI183 VacB KI | Regions of *vacB* in pPI183 | This study |
| pPI183 VacC KI | Regions of *vacC* in pPI183 | This study |
| pKOSG-IBA | Blasticidin<sup>r</sup>, for knock-out | (Wiegand et al., 2011) |
| pJET-VacA-BsR | Regions of *vacA* in pJET with Blasticidin<sup>r</sup> | This study |
| pKOSG-VacB | Regions of *vacB* in pKOSG-IBA | This study |
| pKOSG-VacC | Regions of *vacC* in pKOSG-IBA | This study |
| pKOSG-VacB-VacC | Regions of *vacC* and *vacB* in pKOSG-IBA for double knock-out | This study |
| pJET-VacA-Hyg | Regions of *vacA* in pJET with Hyg<sup>r</sup> for triple knock-out | This study |
| pDM631 PH<sub>CRAC</sub>-GFP |  | (Buckley et al., 2016) |
| pMJC14 GFP-WASH |  | (Park et al., 2013) |
| pDXA-GFP-Rab11b |  | Miriam Essid (TS lab) |

| Mycobacteria plasmids |  |  |
| --- | --- | --- |
| pMSP12::GFP | GFP under control of the MSP promoter, Kan<sup>r</sup> | (Cosma et al., 2004) |
| Bacterial strains |  |  |
| *K. pneumoniae* | Also with expression of GFP | Pierre Cosson (University of Geneva)<br>(Lima et al., 2018) |
| *K. pneumoniae* KP21 |  | (Favre-Bonte et al., 1999) |
| *E. coli* B/r |  | (Gerisch, 1959) |
| *E. coli* DH5α |  |  |
| *P. aeruginosa* DP5 | avirulent isogenic quorum sensing mutants (*ΔtrpD*) | (Alibaud et al., 2008) |
| *M. luteus* |  | (Wilczynska and Fisher, 1994) |
| *B. subtilis* |  | (Ratner and Newell, 1978) |
| *S. aureus* | strain RN6390 | (Wilczynska and Fisher, 1994) |

**Sup. Table 2.**
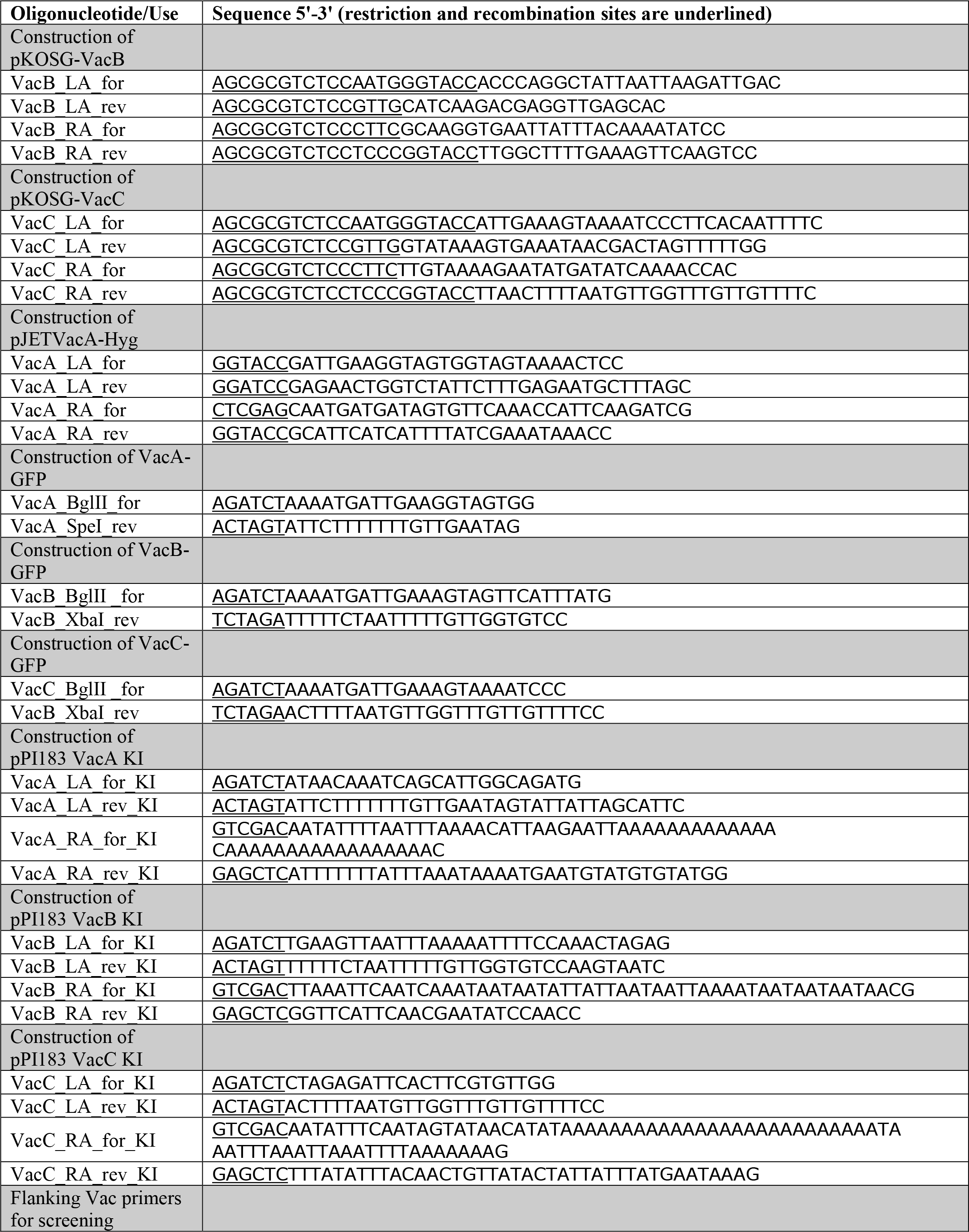
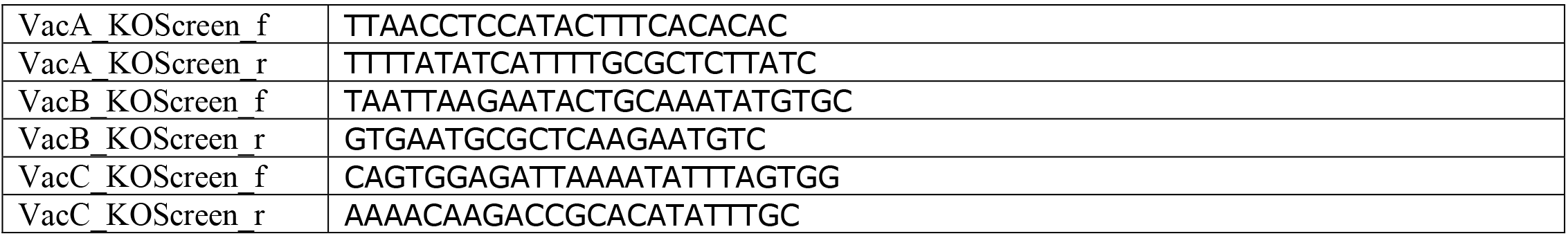
List of oligos used in this study.

**Sup. Table 3.** List of significant (p-value < 0.1) proteins that were over-represented or under-represented at the surface of vacuolin ABC KO compared to wt cells.

| Identified proteins | Accession Number | t-test pvalue | Fold Change | Comments |
| --- | --- | --- | --- | --- |
| Under-represented in ΔABC cells | | | | |
| Protein psiH | Q94494 | 0.099 | 0.6 | TM, glycosylated, PM |
| Protein DD3-3 | Q58A42 | 0.1 | 0.6 | TM, PM |
| Dynamin-like protein A | Q58A42 | 0.087 | 0.6 | cytokinesis |
| Ras-related protein Rab-14 | P36410 | 0.1 | 0.6 |  |
| Penta-EF hand domain-containing protein 1 | Q95YL5 | 0.0048 | 0.6 |  |
| Protein psiL | Q86HT1 | 0.1 | 0.3 | TM, PM |
| Uncharacterized protein | Q75JE7 | 0.095 | 0.3 | TM |
| Uncharacterized protein | Q54X81 | 0.066 | 0.3 | TM |
| Uncharacterized protein | Q55FM2 | 0.095 | 0.3 |  |
| ABC transporter G family member 2 | Q9NGP5 | 0.087 | 0.2 | Several TM, PM |
| Myosin IA heavy chain myoA | P22467 | 0.009 | 0.2 | Cell movement across a surface, extension and retraction of pseudopodia or lamellipodia |
| von Willebrand factor A domain-containing protein | Q54DL7 | 0.039 | 0.2 |  |
| Uncharacterized protein | Q54I84 | 0.049 | 0.1 | EGF-like domain |
| Uncharacterized protein | Q54QY8 | 0.003 | 0.1 | TM |
| Myosin-I heavy chain myoI | Q9U1M8 | 0.0008 | 0.001 | Involved in phagocytosis and adhesion |
| EGF-like domain-containing protein | C7FZX1 | 0.0072 | 0.001 | Pinocytosis, TM, PM |
| Phospholipid-transporting ATPase | Q55E61 | 0.0023 | 0.001 | TM |
| Myosin-K heavy chain myoK | Q9XXV8 | 0.0087 | 0.001 | Involved in phagocytosis, motility, maintenance and dynamics of cell cortex |
| Uncharacterized protein | Q54GQ7 | 0.0008 | 0.001 |  |
| Uncharacterized protein | Q54UB6 | 0.0036 | 0.001 | TM |
| GP63 | Q6TU47 | 0.0036 | 0.001 | TM, cell surface |
| Over-represented in ΔABC cells | | | | |
| Uncharacterized protein | Q55EI7 | 1000 | 0.0001 |  |
| Uncharacterized protein | Q54P41 | 1000 | 0.0003 | Integral TM |
| GTP-binding protein Sar1A | Q559R0 | 4.5 | 0.05 | COPII |
| Uncharacterized protein | Q54DC6 | 4.4 | 0.031 | GTP binding |
| Uncharacterized protein | Q54UX5 | 2.6 | 0.035 |  |
| Luminal-binding protein 2 | Q8T869 | 1.9 | 0.042 | ER |
| ABC transporter G family member 10 | Q54CG0 | 1.8 | 0.046 | Several TM, PM |
| Uncharacterized protein | Q551R4 | 1.7 | 0.085 | Ser/Thr kinase |
| EGF-like domain-containing protein | Q54GV4 | 1.6 | 0.024 | Integral TM |

**Sup. Table 4.** Complete list of proteins identified in surface biotinylation experiments. The list was manually curated and proteins found in the nucleus or mitochondria, or ribosomal proteins were removed.

